# Developmental evidence for parental conflict in driving *Mimulus* species barriers

**DOI:** 10.1101/2022.05.30.494029

**Authors:** Gabrielle D. Sandstedt, Andrea L. Sweigart

## Abstract

The endosperm, a tissue that nourishes the embryo in the seeds of flowering plants, is often disrupted in inviable hybrid seeds between species presumed to have divergent histories of parental conflict. Despite the potential importance of parental conflict in plant speciation, we lack direct evidence of its action in driving species barriers. Here, we performed reciprocal crosses between pairs of three monkeyflower species (*Mimulus caespitosa, M. tilingii*, and *M. guttatus*). The severity of hybrid seed inviability varies among these crosses, which we determined was due to species divergence in effective ploidy. By performing a time series of seed development, we assessed whether regions within the endosperm were potential targets of parental conflict. We found that the chalazal haustorium, a tissue within the endosperm that occurs at the maternal-filial boundary, develops abnormally in hybrid seeds when the paternal parent has the greater effective ploidy. Within these *Mimulus* species, parental conflict might target the chalazal haustorium to control sucrose movement from the maternal parent into the endosperm. Consequently, conflict may be exposed in crosses between species. Our study suggests that parental conflict in the endosperm may function as a driver of speciation by targeting regions and developmental stages critical for resource allocation.

## INTRODUCTION

Identifying the evolutionary drivers of reproductive isolation is critical for understanding the origin of species. This task has been a challenge for intrinsic postzygotic isolation, which arises when hybrids inherit novel combinations of incompatible alleles that cause inviability or sterility (Dobzhansky, 1937; Muller, 1942). Because these incompatible combinations occur uniquely in hybrids and are independent of the environment, there are usually few clues as to why the causal alleles initially increase in frequency and fix within species. In flowering plants, hybrid seed inviability is a common form of postzygotic isolation in which crosses between closely related species produce only flattened, shriveled seeds that fail to germinate (Rebernig *et al*., 2015; Oneal *et al*., 2016; Lafon-Placette *et al*., 2017; Roth *et al*., 2018; Coughlan *et al*., 2020; İltaş *et al*., 2021). Almost invariably, this inviable seed phenotype involves defects in the endosperm (Lafon-Placette & Köhler, 2016), a nutritive tissue that surrounds and feeds the developing embryo. The endosperm is one of two products formed through double fertilization, a key reproductive feature of flowering plants. During this process, one of the haploid pollen sperm cells fuses with the haploid egg cell to form a diploid zygote, while the other fuses with the homodiploid central cell to form a triploid endosperm with a relative contribution of two maternal to one paternal (2m:1p) genomes (Berger, 2003; Berger *et al*., 2008). Given its major role in postzygotic isolation, discovering how the endosperm evolves within and between closely related lineages holds great promise for probing the evolutionary mechanisms of plant speciation.

The first hints that endosperm evolution might drive reproductive barriers came from early crossing studies that showed high rates of seed failure between plants of different ploidies. Many of these studies also reported pronounced reciprocal differences in seed growth and development (Håkasson, 1952; Woodell & Valentine, 1961; Nishiyama & Inomata, 1966). In general, they found that crosses with “maternal excess” – that is, crosses with the higher ploidy plant as the maternal parent – produce smaller seeds than intraploidy crosses that are sometimes inviable. In contrast, “paternal excess” crosses – those with the higher ploidy plant as the pollen donor – generally produce larger seeds, which often abort (Scott *et al*., 1998; Pennington *et al*., 2008; Lu *et al*., 2012). These observations led to the hypothesis that seed failure is caused by a deviation from the usual dosage of 2m:1p genomes in the triploid endosperm (Johnson *et al*., 1980; Lin, 1984; Haig & Westoby, 1989). However, because these same parent-of-origin effects were also discovered in interspecific crosses of the same ploidy (Cooper & Brink, 1942; Stephens, 1949; Nishiyama & Yabuno, 1978), it became clear that disruptions to the 2m:1p ratio can also arise through allelic divergence. Thus, cross compatibility was said to be a function of “effective” ploidy, rather than of absolute genome number (Johnston *et al*., 1980). In this conceptualization, plant species with higher effective ploidies have presumably accumulated genetic variation that mimics the maternal-and paternal-excess effects of higher ploidy plants. Drawing on many of these same classic crossing studies, Haig & Westoby (1991) recognized that this genetic variation must affect functions specific to maternal and paternal genomes and proposed genomic imprinting – parent-specific gene expression – as the underlying mechanism. Indeed, they argued that reciprocal differences in hybrid seed phenotypes between species diverged in effective ploidy are caused by incompatibilities that disrupt imprinted gene regulation.

In addition to offering a molecular mechanism for parent-of-origin effects and hybrid seed inviability, Haig & Westoby (1991) proposed the idea that parental conflict is the evolutionary driver of these phenotypes. Like the mammalian placenta, the angiosperm endosperm plays a critical role in the acquisition and transfer of nutrients to the embryo (Brink & Cooper, 1947). In plant species that receive pollen from more than one donor, the endosperm is predicted to operate as a venue for parental conflict with maternal and paternal genomes evolving different levels of resource acquisition due to their unequal relatedness to offspring (Hamilton, 1964; Haig & Westoby, 1989; Brandvain & Haig, 2005). In a maternal (seed) parent, natural selection should favor gene expression in the endosperm that equalizes nutrient acquisition among all seeds, whereas in a paternal parent (pollen donor), selection should favor gene expression that maximizes resource acquisition in its own offspring at the expense of unrelated seeds (Haig & Westoby, 1989). At a mechanistic level, this scenario is thought to play out through epigenetic modifications during male and female gametogenesis that regulate parent-of-origin biased gene expression in the endosperm (i.e., genomic imprinting; Reik & Walter, 2001; Haig & Westoby, 1991; Kinoshita, 2007; Batista & Köhler, 2020). Within a population, endosperm “balance” should be maintained through coevolution between loci that act to acquire resources from the seed parent and loci that moderate these acquisitive effects; however, species barriers may arise in hybrid genomes formed from species with divergent histories of parental conflict (Haig & Westoby, 1991). Despite the intuitive appeal of this theory, direct evidence for parental conflict in shaping endosperm development and driving barriers between closely related species is fairly limited (but see Coughlan *et al*., 2020). Moreover, because hybrid seed inviability has rarely been investigated in a phylogenic context, we have little understanding of its evolutionary tempo. For example, it is not yet clear how often changes in effective ploidy are tied to shifts in mating system, as might be expected if the evolution of self-fertilization alleviates parental conflict (Brandvain & Haig, 2005), or whether these changes accumulate with genetic distance.

According to the predictions of parental conflict theory, selection in the endosperm should target developmental timepoints or functions that are most important for nutrient uptake (Queller, 1983). Most of what is known about the developmental phenotypes associated with hybrid seed inviability comes from crosses in *Arabidopsis* and other systems with nuclear-type endosperms (so called because the early endosperm forms a syncytium; Bushell *et al*., 2003; Rebernig *et al*., 2015; Floyd & Friedman, 2000), where the timing of cellularization seems to be a major determinant of nutrient acquisition and seed size (Garcia *et al*., 2003; Luo *et al*., 2005; Kang *et al*., 2008; Hehenberger *et al*., 2012). In interploidy crosses in these systems, endosperm cellularization is often precocious when the seed parent has higher ploidy and delayed when the pollen parent has higher ploidy, resulting in smaller or larger seeds, respectively (Scott *et al*., 1998; Pennington *et al*., 2008; Lu *et al*., 2012; Morgan *et al*., 2021). The fact that these same maternal-and paternal-excess effects on cellularization have been observed in crosses between species of the same ploidy in *Arabidopsis* and *Capsella* (Lafon-Placette *et al*., 2017; Rebernig *et al*., 2015; Lafon-Placette *et al*., 2018) has been taken as evidence for parental conflict in nuclear-type endosperms. Although these disruptions in developmental timing are certainly suggestive, few studies of hybrid seed inviability have explicitly investigated resource provisioning functions in distinct regions of the endosperm – especially in systems with non-nuclear modes of endosperm development (i.e., cellular and helobial). In most angiosperms, the endosperm is not a homogeneous structure but rather differentiates into three spatially and functionally distinct domains: the micropylar domain that surrounds the embryo, the chalazal domain that occurs at maternal–filial interface, and the central peripheral domain that makes up the largest portion of the endosperm (Brown *et al*., 2003). Of these domains, the micropylar and chalazal regions appear to be directly involved in nutrient transfer from maternal to filial structures (Baud *et al*., 2005, Morley-Smith *et al*., 2008), making them potential targets of parental conflict and centers for the evolution of reproductive barriers.

Across the wildflower genus *Mimulus*, hybrid seed inviability has evolved repeatedly (Vickery, 1978; Oneal *et al*., 2016; Garner *et al*., 2016; Coughlan *et al*., 2020; Kinser *et al*., 2021; Sandstedt *et al*., 2021), making it an outstanding system for dissecting the developmental and evolutionary mechanisms of this common isolating barrier. In *Mimulus*, the endosperm is of the cellular-type, meaning that cell walls develop following the initial division of the primary endosperm nucleus (Arekal, 1965; Guilford & Fisk, 1952; Oneal *et al*., 2016). After a few rounds of cell division, the three major endosperm domains form (i.e., micropylar, chalazal, and central-peripheral endosperm), with the micropylar and chalazal regions giving rise to separate haustoria that likely act as channels for nutrient transfer between the maternal plant and developing seed (Nguyen *et al*., 2000; Mikesell, 1990). The chalazal haustorium is ephemeral, composed of two cells extending from the ovule toward the micropylar domain that typically degenerates when the embryo is near a globular stage (Arekal, 1965; Guilford & Fisk, 1952; Oneal *et al*., 2016). On the opposite end of the seed, the two cells of the micropylar haustorium appear to penetrate the integuments (i.e., precursors of the seed coat) and degenerate when the embryo is nearly fully developed (Arekal, 1965). Given their invasion of neighboring tissues to funnel nutrients to the developing embryo, we might expect defects in the haustoria of hybrid seeds if *Mimulus* species have diverged in their levels of parental conflict. Such phenotypes have been noted before in chalazal structures of interploidy crosses in *A. thaliana* (Scott *et al*., 1998), but they have not been described in a conflict scenario between species of the same ploidy.

In this study, we investigate the developmental phenotypes associated with hybrid seed inviability among three closely related, diploid *Mimulus* species with a nested pattern of relatedness: *M. caespitosa* and *M. tilingii* shared a common ancestor ∼382 kya, and *M. guttatus* diverged from the other two ∼674 kya (Sandstedt *et al*., 2021). Populations of *M. caespitosa* and *M. tilingii* occur exclusively at high elevations and appear to be mostly allopatric, with *M. caespitosa* restricted to Washington state and *M. tilingii* mostly known from alpine areas of Oregon and California. *M. guttatus* occupies a more diverse range in western North America, sometimes overlapping with populations of *M. caespitosa* and *M. tilingii* (Nesom, 2012; Coughlan *et. al*., 2021). Previously, we showed that crosses between *M. caespitosa* and *M. tilingii* result in severe hybrid seed inviability – but only when *M. tilingii* is the paternal parent (crosses in the reciprocal direction produce mostly viable seeds, Sandstedt *et al*., 2021). Hybrid seed inviability is even stronger between the more distantly related *M. tilingii* and *M. guttatus*, which produce very few (< 1%) viable seeds in either direction of the cross (Vickery, 1978; Garner *et al*., 2016). Despite this apparent similarity between reciprocal crosses of *M. tilingii* and *M. guttatus*, most of the underlying genetic loci affect seed viability only through the maternal or paternal parent (Garner *et al*., 2016). These parent-of-origin effects on seed viability and genetic loci strongly point to a role for the endosperm, but its involvement has not yet been directly tested.

Here, we leverage this closely related trio of *Mimulus* species to investigate whether parental conflict is an important driver of hybrid seed inviability. First, we explore the severity of hybrid seed inviability in each of the three species pairs and determine whether the endosperm is involved. Second, we investigate divergence in effective ploidy among the three *Mimulus* species. For each species pair, we ask whether increasing the ploidy of one species can “balance” the genetic contribution of the other and rescue hybrid seed inviability. We use this genome doubling approach to establish hierarchical relationships in effective ploidy among the three species and determine how it scales with genetic distance. Finally, we investigate the role of parental conflict in shaping this hierarchy and driving species barriers. We perform detailed developmental analyses of pure species and hybrid seeds, asking whether developmental phenotypes linked to resource acquisition appear particularly affected by divergence in effective ploidy. Together, our results provide strong evidence for parental conflict as a driver of reproductive isolation in this group of *Mimulus* species.

## MATERIALS AND METHODS

### Generation of Plant Material

Here, we used one inbred line (formed from ≥8 generations of self-fertilization) for each focal species (*M. caespitosa, M. tilingii*, and *M. guttatus*). The same inbred lines were used in previous studies of hybrid seed inviability in *M. tilingii* and *M. guttatus* (Garner *et al*., 2016) and *M. caespitosa* (Sandstedt *et al*., 2021). The *M. caespitosa* inbred line, TWN36, originates from a high-alpine population at 1594m in Twin Lakes, WA. The *M. tilingii* inbred line, LVR1, is derived from a population at 2751m in Yosemite Park, CA. The *M. guttatus* inbred line, DUN10, originates from a population in the Oregon Dunes National Recreation Area.

In this study, we considered three intraspecific crosses (CxC, TxT, and GxG, where C = *M. caespitosa*, T = *M. tilingii*, and G = *M. guttatus*) and six interspecific crosses (CxT, TxC, TxG, GxT, CxG, GxC; maternal parent is always listed first). To generate diploid, experimental plants, we sowed 20-30 seeds for each inbred line on damp paper towels in petri dishes sealed with parafilm and cold-stratified them for 7 days to disrupt seed dormancy. After cold stratification, we transferred petri dishes to a growth chamber with 16-h days at 23°C and 8-h nights at 16°C. We transplanted seedlings into 3.5” pots with moist Fafard 4P growing mix (Sun Gro Horticulture, Agawam, MA) and placed the pots in the same growth chamber. Once plants began flowering, we randomly crossed within and between individuals (total plants: C = 19, T = 19, G = 15). For all crosses, we emasculated the maternal plant 1-3 days prior to each cross to prevent contamination from self-pollination.

To investigate species divergence in effective ploidy, we performed several interspecific, interploidy crosses: C_4n_xT, TxC_4n_; T_4n_xG, GxT_4n_; C_4n_xG, GxC_4n_ (4n subscript indicates tetraploid). To generate synthetic tetraploid individuals, we treated 100-200 seeds of TWN36 and LVR1 with 0.1% or 0.2% colchicine and stored them in the dark for 24 hours 16 hours at 23°C and 8 hours at 16°C. The next day, we planted seeds onto Fafard 4P potting soil using a pipette and placed pots inside the growth chamber under typical light and temperature conditions (16-h days at 23°C and 8-h nights at 16°C). Once seeds germinated, we transplanted seedlings into 2.5” pots. After sufficient growth, we prepared samples for flow cytometry using a protocol adapted from Lu *et al*., 2017. Briefly, we extracted nuclei from one colchicine-treated sample and an internal control (2n *Mimulus* or *Arabidopsis thaliana*, Col-0) together in a single well. To extract nuclei, we chopped 100mg of leaf tissue (50mg colchicine-treated sample and 50 mg internal control) in 1mL of a pre-chilled lysis buffer (15mM Tris-HCl pH 7.5, 20mM NaCl, 80mM KCl, 0.5mM spermine, 5mM 2-ME, 0.2% TritonX-100). We stained nuclei with 4,6-Diamidino-2-phenylindole (DAPI), filtered nuclei for debris using a 40um Flowmi™ cell strainer, and aliquoted nuclei into a single well of a 96-well polypropylene plate. We assessed ploidy of each sample using a CytoFLEX (Beckman Coulter Life Sciences) flow cytometer. We calculated total DNA content using the following equation:

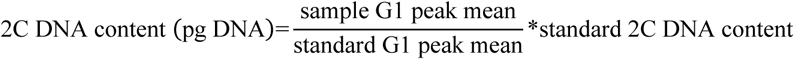

We generated three synthetic polyploids for TWN36 and six for LVR1. For each synthetic polyploid, 2C DNA content was nearly doubled compared to corresponding diploid lines (TWN36, 2C = 1.38 pg; TWN36_4n_, 2C = 2.69 ±0.09 pg; LVR1, 2C = 1.26 pg; LVR1_4n_, 2C = 2.64 ±0.05 pg). In some cases, we discovered that plants initially identified as tetraploid via flow cytometry were actually mixoploids. To ensure the crosses we performed were indeed interploidy, we determined the ploidy of the resulting progeny. From each interploidy cross, we planted 5-10 seeds per fruit, isolated nuclei from the resulting plants, and assessed 2C content using a flow cytometer for a few offspring as described above (3n TWN36_4n_xLVR1 = 1.92 ±0.04, 3n LVR1xTWN36, 2C = 1.88 ±0.01 pg; 3n LVR1_4n_xDUN10, 2C = 1.95 ±0.04 pg; 3n DUN10xLVR1_4n_, 2C = 1.81 ±0.01 pg). We included data from interploidy crosses only when their progenies were confirmed to be triploids, or, in the case of 4n *M. caespitosa*, if we were using a confirmed stable polyploid line (i.e., self-fertilized at least one generation with polyploidy confirmed in the progeny).

### Measuring seed size and seed viability

To measure seed size, we collected three replicate fruits per cross, with each fruit collected from a distinct plant. We imaged 50 seeds per fruit under a dissecting scope, for a total of 150 seeds per cross (except for one CxG fruit for which only 35 seeds were measured for a total of 135 seeds). Seed area was measured using ImageJ (Rasband, 1997).

Using these same fruits, as well as fruits from interploidy crosses (2-5 fruits/cross, at least two fruits per cross from a distinct plant), we assessed seed viability using two different methods. First, we performed visual assessments of mature seeds, looking for irregular phenotypes (shriveled, wrinkled, or flat) known to be associated with hybrid seed inviability in *Mimulus* (Garner *et al*., 2016, Oneal *et al*., 2016, Coughlan *et al*., 2020, Sandstedt *et al*., 2021). We scored the number of seeds that appeared round and plump (*i*.*e*., fully-developed) versus irregularly shaped (i.e., under-developed). Second, we performed Tetrazolium assays to assess seed viability on a subset of these same seeds (∼100 seeds per fruit). For fruits generated from interploidy crosses and fruits that produced <100 seeds, we stained 32-63 seeds. We immersed seeds in a scarification solution (83.3% water, 16.6% commercial bleach, and 0.1% Triton X-100) and placed them on a shaker for 15 minutes. After scarification, we washed seeds five times with water and incubated seeds with 1% Tetrazolium at 30°C. Two days later, we scored the number of seeds that stained dark red (viable) versus pink or white (inviable).

### Seed viability rescues

To assess whether aberrant endosperm development contributes to seed defects in interspecific crosses, we attempted to rescue seed viability with a sucrose-rich medium. We collected three fruits 8 to 12 days after pollination (DAP) from each intra-and interspecific cross (not including interploidy crosses), with each fruit collected from a distinct plant. Of the three fruits per cross, at least one fruit was collected 8 DAP (to maximize the chance of rescue). On average, we dissected 40 whole immature seeds per fruit (range = 25-57) and placed them on petri dishes with MS media containing 4% sucrose. We sealed petri dishes with parafilm and placed them at 23°C with constant light for 14 days before scoring germination.

### Visualizing parent-of-origin effects during seed development

To compare trajectories of seed development, we performed intra-and interspecific crosses, and we collected fruits 3, 4, 5, 6, 8, and 10 DAP. For consistency, we performed crosses and collected fruits at the same time of day.

To visualize early seed development, we collected fruits 3 and 4 DAP (*N* = 1 to 2 fruits per DAP per cross) and prepared them for clearing with Hoyer’s solution. We placed developing fruits in a 9 EtOH: 1 acetic acid fixative overnight. The following day, we washed fruits twice in 90% EtOH for 30 min per wash. We dissected immature seeds directly from the fruit onto a microscope slide with 100uL of 3 parts Hoyer’s solution (70% chloral hydrate, 4% glycerol, 5% gum arabic): 1 part 10% Gum Arabic and sealed the slide with a glass cover slip. We stored the microscope slides containing cleared, immature seeds at 4°C overnight. The next day, we imaged slides using the differential interference contrast (DIC) setting with the 20x objective on a Leica DMRB microscope. For each fruit, we scored the number of developing seeds with and without an intact chalazal haustorium (15-56 seeds per fruit; 32-111 seeds per cross per DAP); only seeds with visible embryos were scored. Additionally, we imaged an average of 11 seeds per fruit (3-15 seeds per fruit, 10-27 seeds per cross per DAP) to assess size differences in the endosperm and chalazal haustorium at 3 and 4 DAP. For the interploidy T_4n_xG cross, we imaged on average 18 seeds per fruit (14-26 seeds per fruit, 29-40 seeds per cross per DAP). We outlined and measured the endosperm in all seeds and the chalazal haustorium when present using ImageJ (Rasband, 1997). Because the chalazal haustorium was not present for all imaged seed, sample sizes for its measurements were lower. We selected and measured images that represented typical seed development at each time point.

We defined the chalazal haustorium as two uninucleate cells that, together, form a continuous structure that penetrates toward the ovule hypostase cells (a group of tightly packed cells at the base of the ovule). To measure the chalazal haustorium, we began the outline near the epidermis of the seed (not including the hypostase cells) and extending it toward the micropylar region following Guilford & Fisk, 1952 (see their Figure 27). In addition, when measuring the endosperm, we started the outline at the same position near the epidermis of the ovule and extended it toward the opening of the micropylar haustorium.

To visualize later seed development (after 4 DAP when the seed coat is too thick to clear with Hoyer’s solution), we collected whole fruits at 5, 6, 8, and 10 DAP and stored them in a Formaldehyde Alcohol Acetic Acid fixative (10%:50%:5% + 35% water) for a minimum of 48 hours. After fixation, we dehydrated developing fruits with increasing concentrations of Tert

Butyl Alcohol. Next, we washed fruits three times for two hours each with paraffin wax at 65°C before embedding them into a wax block. We sectioned wax blocks containing whole fruits into ribbons using a LIPSHAW Rotary Microtome (Model 45). Fruits collected at 5 and 6 DAP were sectioned into 12-um ribbons for better visualization of micropylar and chalazal domains, and fruits collected at 8 and 12 DAP were sectioned into 8-um ribbons. Next, we gently placed ribbons in a warm (∼40°C) water bath and positioned them onto a microscope slide. We placed slides on a slide warmer overnight to adhere sections completely to the glass. In a staining series, we first used Xylene as a clearing agent and performed several washes with increasing concentrations of EtOH to effectively stain nuclei and cytoplasm (1% Safranin-O and 0.5% Fast Green, respectively). We further washed stained slides with EtOH and finished the series with Xylene. We sealed slides with a glass coverslip using Acrytol as the mounting medium.

We visualized slides using a Zeiss Axioskop 2 microscope with a 10x objective. For each fruit, we imaged at least 10 seeds with a developing embryo per fruit (except for severe embryo-lethal crosses: 10 DAP TxG, 8 seeds imaged; 10 DAP CxG, 1 seed imaged). We imaged at least five consecutive sections of each seed through the embryo. For all seeds imaged at 5 and 6 DAP, we scored the presence of the chalazal haustorium. Additionally, we categorized embryo development at 6, 8, and 10 DAP into four different stages: before globular to globular, late-globular to transition, early-heart to late-heart, and torpedo.

### Data Analysis

We performed several statistical analyses to determine the effect of each cross on seed area, seed viability, germination success on sucrose, and area of the endosperm filled by the chalazal haustorium. For each seed phenotype, we used the R software package (Bates *et al*., 2007) to generate a linear model, linear mixed model, or a generalized linear mixed model. Details of each model are described in Methods **S1**.

## RESULTS

### *A central role for the endosperm in* Mimulus *hybrid seed inviability*

Hybrid seed inviability is an exceptionally strong isolating barrier in crosses between *Mimulus guttatus, M. tilingii*, and *M. caespitosa* (Figs. **1a**, **S1**, Tables **S1, S2**). Consistent with our earlier work (Garner *et al*., 2016), *M. guttatus* and *M. tilingii* produced almost exclusively inviable F1 hybrid seeds in both directions of the cross. We found this same result in crosses between *M. guttatus* and *M. caespitosa*. On the other hand, as we have shown previously (Sandstedt *et al*., 2021), F1 hybrid seed inviability between the more closely related *M. tilingii* and *M. caespitosa* occurs in only one direction of the cross.

**Fig. 1.**
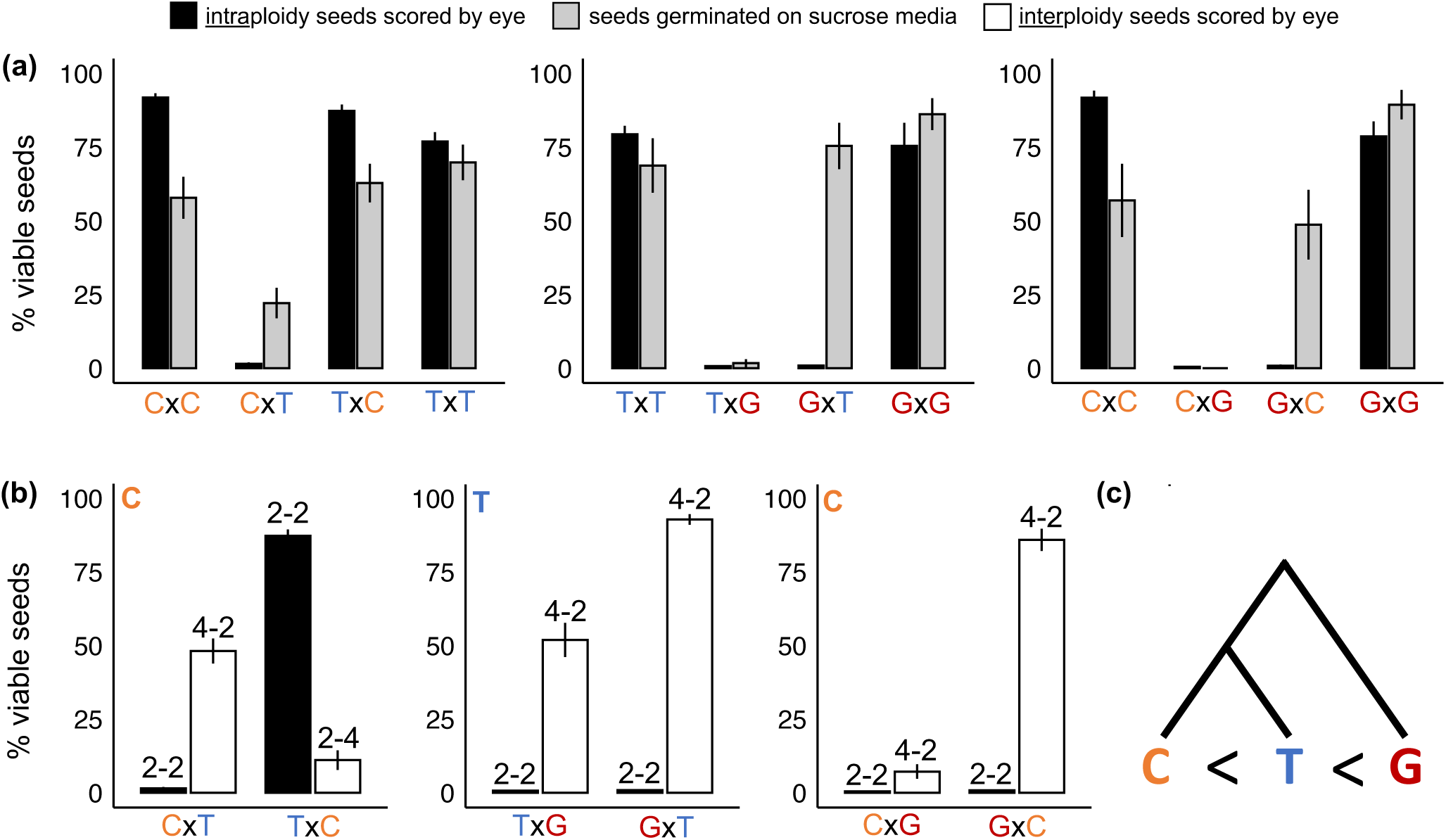
Percentage of viable seeds from intra- and interspecific crosses among *M. caespitosa* (C), *M. tilingii* (T), and *M. guttatus* (G). The first letter of each cross indicates the maternal species. Least squares means (lsmeans) given with +/-SE. Models were generated separately, comparing reciprocal interspecific crosses and their corresponding intraspecific crosses for any given seed viability test *(i*.*e*., fully-developed seeds scored by eye and seeds germinated on sucrose media). Note that reciprocal <u>inter</u>ploidy crosses were included for each model of seed viability scored by eye. (**a)** Percent seeds per fruit that appeared fully-developed (black bars) and percent seeds rescued by a sucrose medium (gray bars). (**b)** Percent seeds per fruit that appeared fully-developed from interspecific <u>intra</u>ploidy (black bars) and <u>inter</u>ploidy (white bars) crosses. The numbers above the bars indicate interspecific crosses between the same (“2-2”) and different (“4-2”, “2-4”) ploidy levels with the maternal parent’s ploidy listed first. The letter in the top left corner of each plot indicates the tetraploid species in the interploidy crosses. (**c)** Simplified phylogenetic tree (modified from Sandstedt *et al*. 2021) with effective ploidy relationships among the three species: *M. caespitosa* is the lowest, *M. tilingii* is intermediate, and *M. guttatus* is the highest.

To investigate endosperm involvement in *Mimulus* hybrid seed failure, we attempted to rescue inviable seeds by plating them on a nutritive, sucrose medium. Even when reciprocal F1 hybrid seeds appear similar in terms of morphology (i.e., flat and shriveled), supplying them with sucrose revealed clear reciprocal differences in viability (Fig. **1a**, Table **S3**). With *M. guttatus* as the maternal parent, F1 hybrid seeds from crosses with *M. tilingii* or *M. caespitosa* germinate on sucrose at rates similar to seeds from parental crosses. In contrast, F1 hybrid seeds with *M. guttatus* as the paternal parent remain almost completely inviable even when supplied with sucrose. This result might indicate that hybrid seed inviability is independent of the endosperm, or that the endosperm defect is so severe that embryo development is irreversibly damaged. In any case, these stark reciprocal differences in F1 hybrid seed inviability – with and without sucrose – point to a central role for the endosperm in reproductive isolation between these *Mimulus* species.

### *Divergence in effective ploidy among* Mimulus *species*

To investigate differences in effective ploidy among this trio of *Mimulus* species, we performed a series of interploidy crosses, testing whether artificially doubling the genome content of one parent could alleviate hybrid seed inviability. Using this approach, we discovered additional support for endosperm-based barriers and determined the rank order of effective ploidy among the three *Mimulus* species (Fig. **1b**, Tables **S1, S2**). Consistent with *M. caespitosa* having the lowest effective ploidy, doubling its genome greatly improves hybrid seed viability in crosses with *M. tilingii* – but only when *M. caespitosa* acts as the seed parent. In the reciprocal direction, which normally produces viable seeds (Fig. **1a**), 4n *M. caespitosa* pollen donors actually induce seed inviability. These results illustrate that divergence in effective ploidy can cause distinct effects through the two parental genomes: paternal excess from *M. tilingii* is severe enough to cause seed inviability, whereas maternal excess is sufficiently modest that increasing paternal dosage from *M. caespitosa* overcompensates for its effects. Along this continuum of effective ploidy, *M. guttatus* has diverged even further: 4n *M. caespitosa* restores F1 hybrid seed viability only minimally when it acts as the seed parent in crosses with this species, indicating severe paternal excess stemming from *M. guttatus*. On the other hand, maternal-excess inviability from *M. guttatus* is not as debilitating: GxC F1 hybrid seeds are completely rescued by doubling the genome content of *M. caespitosa*. Among the three species, *M. tilingii* has an effective ploidy that is intermediate to the other two, with crosses between 4n *M. tilingii* and *M. guttatus* largely or completely restoring hybrid seed inviability. Taken together, these results demonstrate clear differences in effective ploidy: *M. guttatus* has the highest, *M. tilingii* is intermediate, and *M. caespitosa* has the lowest (Fig. **1c**).

### *Developmental phenotypes in* Mimulus *hybrids implicate parental conflict*

To investigate whether parental conflict is the evolutionary force driving these changes in effective ploidy, our next step was to take a closer look at parent-of-origin seed phenotypes. As a first pass, we examined reciprocal differences in F1 hybrid seed size for each species pair, reasoning that maternal-excess crosses might show signs of undergrowth and paternal-excess crosses might show signs of overgrowth. Contrary to this expectation, hybrid seeds are almost always smaller than pure species seeds (except for CxT, which are the same size) and reciprocal differences are subtle or absent (Fig. **S2**, Table **S4**). However, because mature hybrid seed size depends on a multitude of developmental processes, including embryo growth and early seed abortion, it might not reflect parent-of-origin phenotypes operating during development.

Indeed, despite superficial similarities in seed size, we observed dramatic differences in the underlying development of all reciprocal pairs of F1 hybrid seeds. In early seed development, we observed overgrowth of the chalazal haustorium in all paternal-excess crosses (CxT, TxG, CxG in Figs. **2a**, **S3, S4**, Table **S5**). Whereas during normal seed development (i.e., in the progeny of intraspecific crosses CxC, TxT, and GxG), the chalazal haustorium decreases in size early (3-4 DAP) and degenerates completely by 5 DAP, it occupies a significantly larger proportion of the endosperm in paternal-excess crosses and is maintained much longer (Figs. **3**, **4**, **S3**). In the paternal-excess cross between *M. caespitosa* and *M. tilingii*, the volume of endosperm devoted to the chalazal haustorium at 4 DAP is nearly twice that of viable seeds (compare CxT to CxC, TxT, and TxC, Figs. **2a**, **3**, **S3**, Table **S5**) and chalazal structures are maintained until 6 DAP (Fig. **4**, **S5**). Developmental irregularities in chalazal haustoria are even clearer in paternal-excess crosses involving *M. guttatus*, the species with the largest effective ploidy: in TxG and CxG F1 hybrid seeds, the proportion of the endosperm filled by the chalazal haustorium is ∼3-4x greater than in the seeds of reciprocal and intraspecific crosses, and haustoria persist through 6 DAP (Figs. **2a**, **3**, **4**, **S5**, Table **S5**). Remarkably, this developmental defect is almost completely rescued by increasing maternal dosage. Indeed, the volume of endosperm filled by chalazal haustoria is greatly reduced in 4n *M. tilingii* x *M. guttatus* hybrids (Figs. **2b**, **3**, Table **S5**) and haustoria are almost entirely degenerated by 4 DAP (Fig. **4**).

**Fig. 2.**
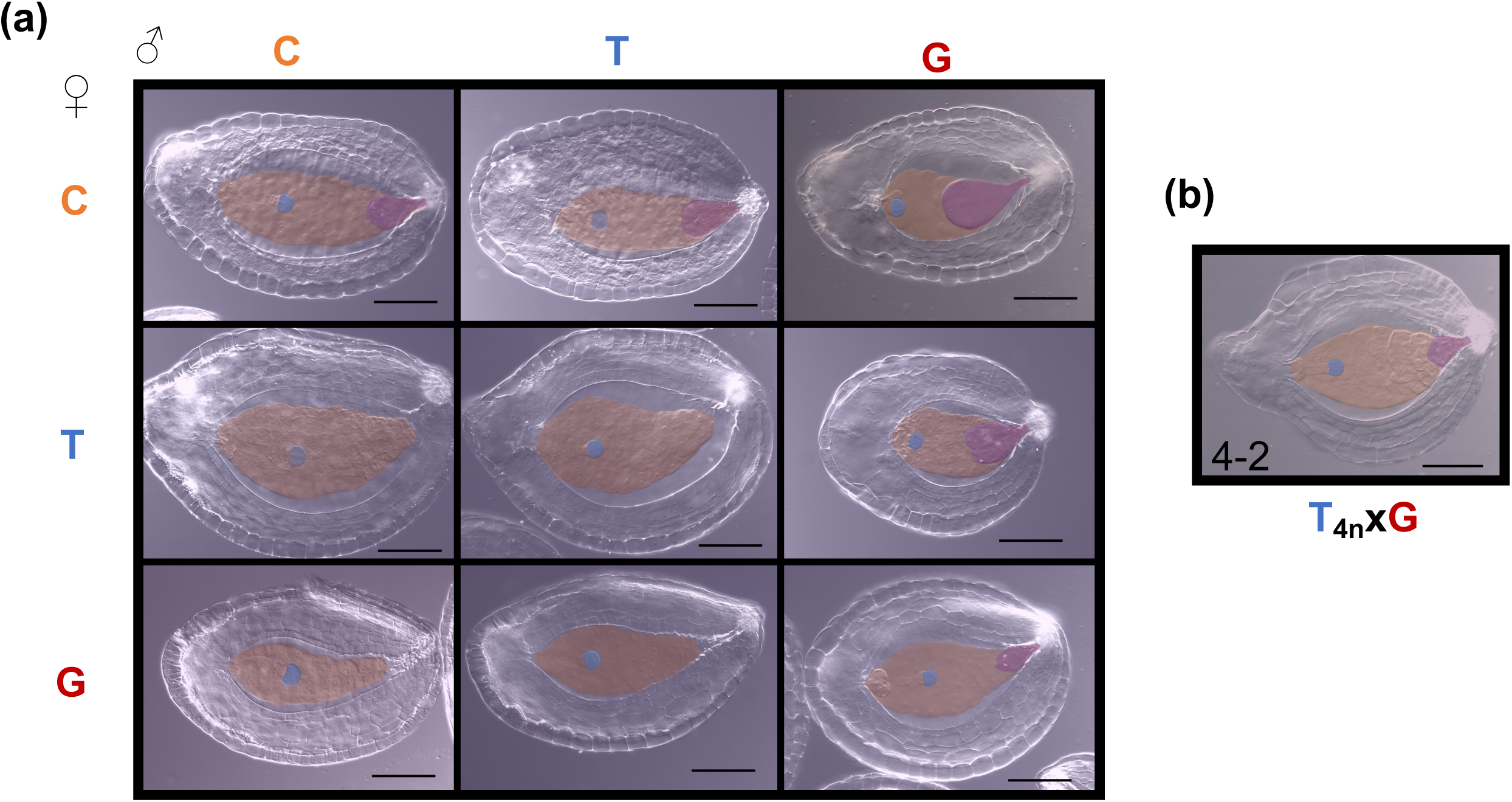
Developing seeds four days after pollination (DAP) in crosses among *M. caespitosa* (C), *M. tilingii* (T), and *M. guttatus* (G). Developing seeds were cleared with Hoyer’s solution. Structures were outlined and artificially shaded: blue shading represents embryo, orange shading represents endosperm region, and purple shading represents chalazal haustorium. Scale bar is 0.1mm. (**a)** Seeds 4 DAP of intra- and interspecific crosses. Maternal parent is listed along the left side, and paternal parent is listed along the top. Along the diagonal are the intraspecific crosses (CxC, TxT, and GxG), below diagonal are maternal-excess crosses (CxT, GxT, and GxC), and above diagonal are paternal-excess crosses (CxT, TxG, and CxG). (**b)** Representative seed of interploidy cross at 4 DAP. In the bottom left corner, “4-2” represents that the cross was between two ploidy levels, with the tetraploid maternal parent ploidy listed first. In addition, the “4n” subscript in T_4n_xG indicates that the maternal *M. tilingii* parent is a synthetic tetraploid.

**Fig. 3.**
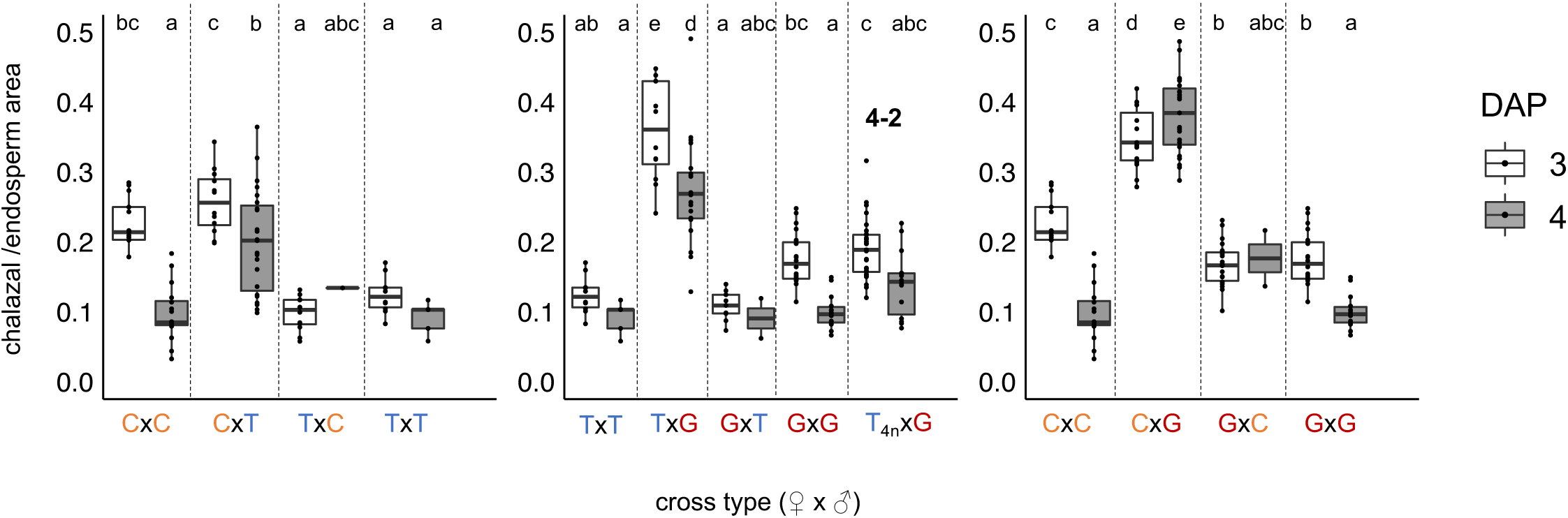
Proportion of endosperm filled by a chalazal haustorium at 3 and 4 days after pollination in intra- and interspecific crosses among *M. caespitosa* (C), *M. tilingii* (T), and *M. guttatus* (G). The first letter of each cross indicates the maternal species. In the T_4n_xG cross, “4n” subscript indicates a synthetic tetraploid *M. tilingii* maternal parent. Further, “4-2” denotes that the cross was performed between two ploidy levels – tetraploid maternal parent and diploid paternal parent. Different letters above boxes indicate significant differences in least squares means among crosses (*P*<0.05) determined by a post hoc Tukey method. Analyses were performed separately, comparing reciprocal interspecific and corresponding intraspecific crosses, except for crosses between *M. tilingii* and *M. guttatus* that include comparisons with the T_4n_xG cross.

**Fig. 4.**
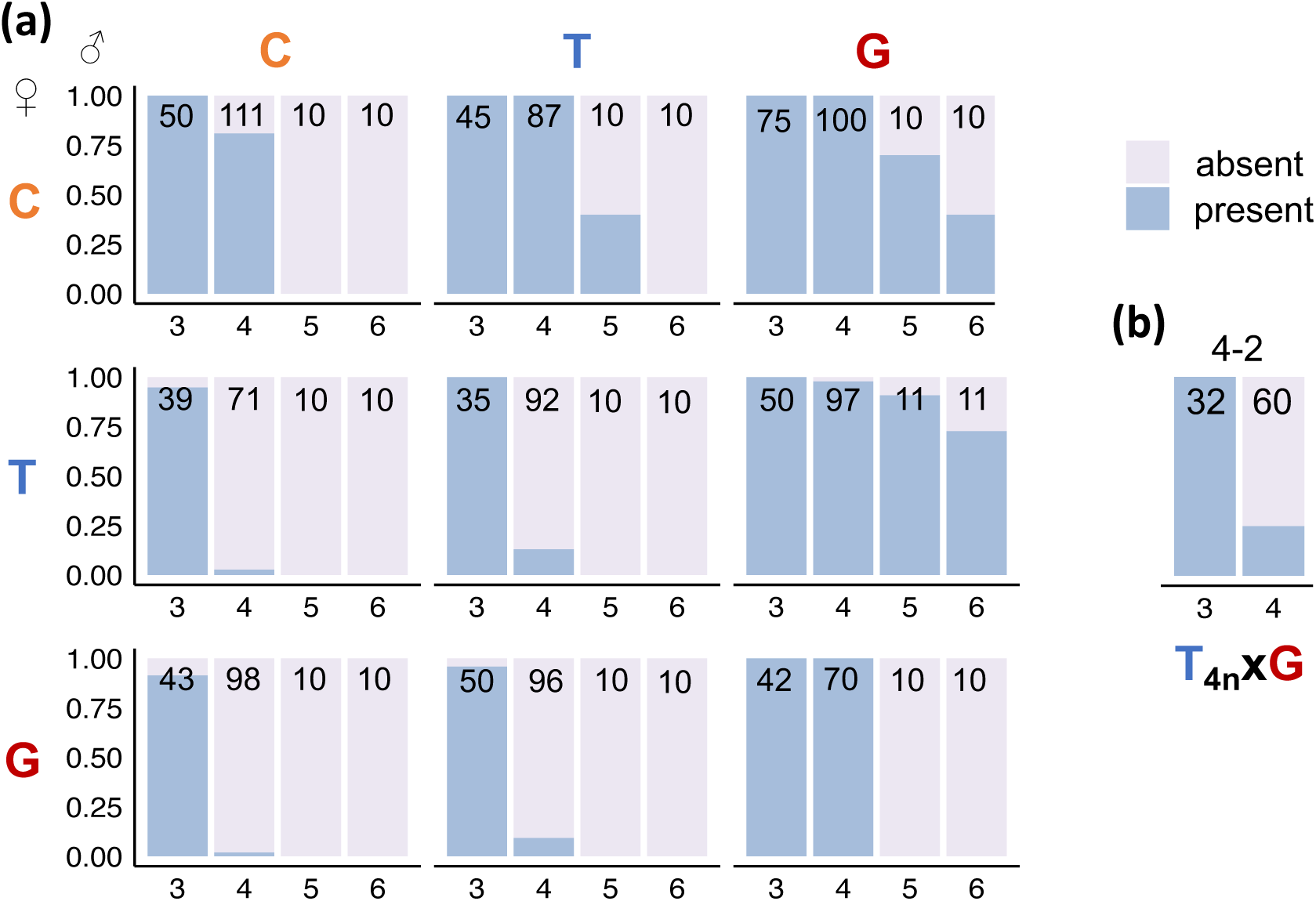
Proportion of developing seeds with a chalazal haustorium (3, 4, 5, and 6 days after pollination) from intra- and interspecific crosses among *M. caespitosa* (C), *M. tilingii* (T), and *M. guttatus* (G). Numbers in bars represent the total number of developing seeds scored for a chalazal haustorium, with seeds dissected from 1-2 fruits per cross type per DAP. Seeds were only scored and imaged if they contained a visible embryo. The blue color represents the proportion of seeds with a chalazal haustorium, and the purple color represents the proportion of seeds without a chalazal haustorium. At days 3 and 4, chalazal haustorium presence/absence was scored after dissecting developing seeds from whole ovules and clearing them with Hoyer’s solution. At days 5 and 6, this phenotype was scored from whole fruit, histological sections. **(a)** Along the diagonal are the intraspecific crosses (CxC, TxT, and GxG), below diagonal are maternal-excess crosses (CxT, GxT, and GxC; maternal parent always listed first), and above diagonal are paternal-excess crosses (CxT, TxG, and CxG). **(b)** In the T_4n_xG cross type, “4n” subscript denotes synthetic tetraploid *M. tilingii* maternal parent. The “4-2” above the bars further represents the cross between two ploidy levels, with a tetraploid maternal parent and diploid paternal parent.

Parent-of-origin effects in the endosperm become even more apparent at later stages of development. At 6 DAP, the embryo of most pure species seeds is at the globular-to-transition-stage and is surrounded by a cellularized endosperm with cells that appear largely empty (Figs. **5a**, **6**, **S5**). By 8 DAP, the centrally-located endosperm cells of these normally developing seeds begin to break down, while the peripheral endosperm lining the seed coat differentiates into cytoplasmically dense, starch-filled cells (Figs **5b**, **S5**). However, in maternal-excess crosses, especially those with *M. guttatus* as the seed parent, these differentiated endosperm cells appear earlier (6 DAP) and are tightly packed into a much smaller area, leaving little space for embryo progression. As a result, embryos of seeds from *M. guttatus* maternal-excess crosses fail to transition from the heart to the torpedo stage (TxG, CxG in Figs. **5**, **6**, **S5**). Paternal-excess crosses, on the other hand, produce hybrid seeds with delayed endosperm differentiation accompanied by stymied embryo development (CxT in Figs. **5**, **6**, **S5**). In the most severe paternal-excess crosses (involving *M. guttatus* as the pollen parent), the endosperm cells of hybrid seeds fail to differentiate at all and persist as large, empty cells unable to support embryo development past the globular stage (TxG and CxG in Figs. **5**, **6**, **S5**).

**Fig. 5.**
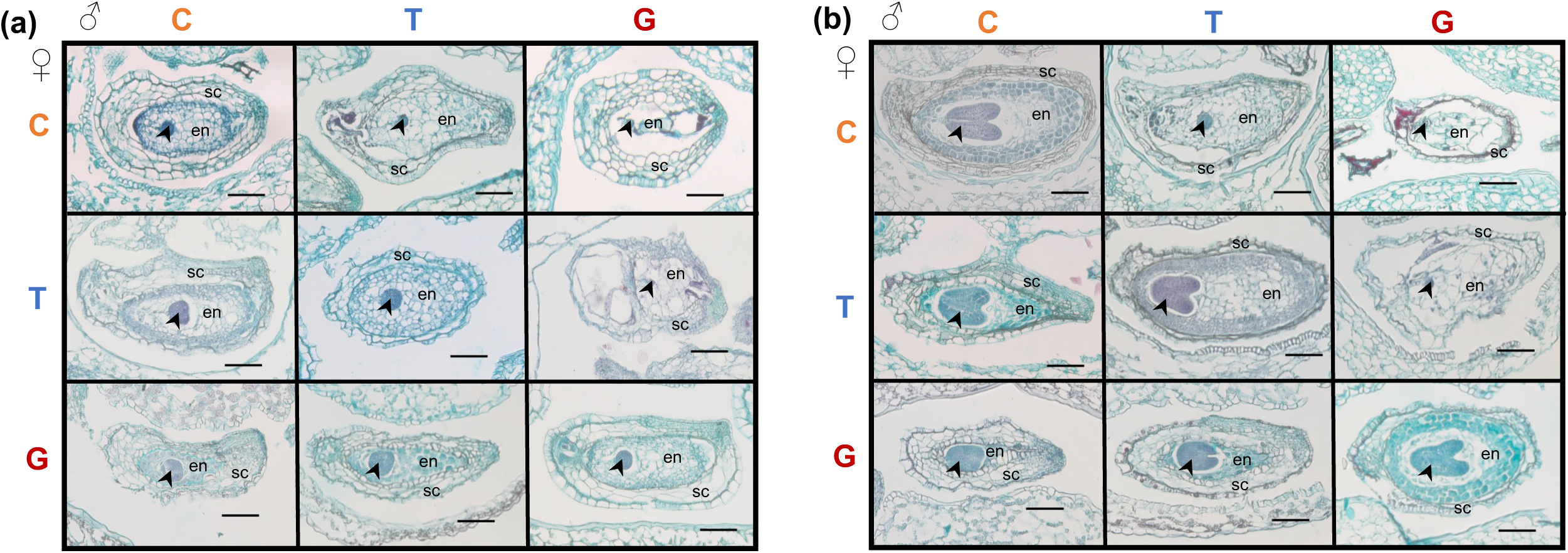
Histological sections of whole fruits from intra- and interspecific crosses among *M. caespitosa* (C), *M. tilingii* (T), and *M. guttatus* (G). Maternal parent is listed along the left side, and paternal parent is listed along the top. Along the diagonal are the intraspecific crosses (CxC, TxT, and GxG), below diagonal are maternal-excess crosses (CxT, GxT, and GxC; maternal parent always listed first), and above diagonal are paternal-excess crosses (CxT, TxG, and CxG). Arrowhead = embryo, en = endosperm, sc = seed coat. Scale bar is 0.1mm. **(a)** 6 DAP. Intraspecific and paternal-excess endosperms are mostly composed of large empty cells, whereas maternal-excess crosses (especially GxT and GxC) develop endosperms that are small and composed of darkly stained, dense cells. **(b)** 8 DAP. Intraspecific endosperm cells begin to differentiate into cytoplasmically dense, starch-filled cells along the peripheral region near the seed coat. However, in GxT and GxC crosses, the whole endosperm is composed of these dense cell types, and the endosperm remains very small and compact. Paternal-excess endosperms appear abnormal and do not show evidence of cell differentiation by 8 DAP.

**Fig. 6.**
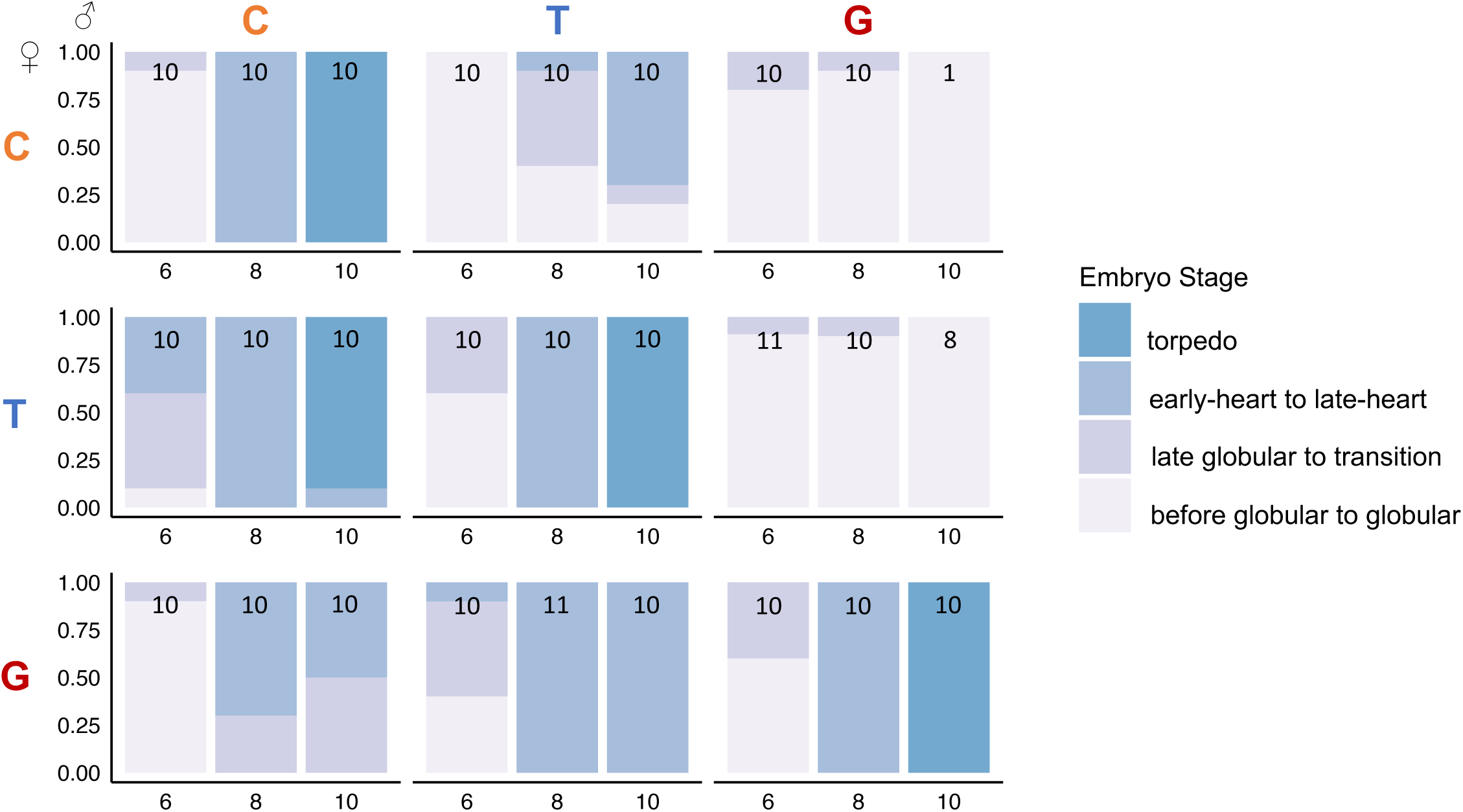
Proportion of embryos at a particular developmental stage at different time points (6, 8, 10 DAP) in intra- and interspecific crosses among *M. caespitosa* (C), *M. tilingii* (T), and *M. guttatus* (G). Numbers in bars represent the total number of embryos scored per cross type, where less than 10 embryo suggests severe embryo lethality for a particular cross type. Colors in each bar represents stage of embryo development: lighter purple color represents early to globular embryos, the darker purple color represents late globular to transition embryos, the lighter blue color represents early to late heart stage embryos, and the darker blue color represents torpedo embryos. Stages of embryo development determined from whole fruit histological sections. Along the diagonal are the intraspecific crosses (CxC, TxT, and GxG), below diagonal are maternal-excess crosses (CxT, GxT, and GxC; maternal parent always listed first), and above diagonal are paternal-excess crosses (CxT, TxG, and CxG).

## DISCUSSION

Identifying the evolutionary drivers of reproductive isolation is a central goal of speciation but remains a formidable challenge, especially for intrinsic postzygotic barriers. Our study provides some of the strongest empirical evidence to date for parental conflict as potent force in the evolution of hybrid seed inviability. Here, we determined that three closely related *Mimulus* species differ in effective ploidy and that crosses between any species pair results in nearly complete reproductive isolation. By performing a detailed time series of normal and F1 hybrid seed development, we uncovered prominent phenotypes with parent-of-origin effects that strongly implicate parental conflict in divergence among *M. caespitosa, M. tilingii*, and *M. guttatus*. This study is one of the first to detail the disruption of nutrient acquiring tissues within the endosperm from hybridizations between species of the same ploidy.

Theory predicts parental conflict should specifically target the developmental structures and processes most closely connected to offspring nutrient acquisition (Queller, 1983; Haig & Westoby, 1989). Support for this idea from well-studied systems like *Arabidopsis* and *Capsella* – both with the nuclear mode of endosperm development – has centered around the timing of endosperm cellularization. In maternal-excess crosses, precocious cellularization leads to reduced nuclear proliferation and seed size, whereas in paternal-excess crosses, delayed cellularization results in nuclei over-proliferation and larger seeds (Scott *et al*., 1998; Pennington *et al*., 2008; Rebernig *et al*., 2015; Lafon-Placette *et al*., 2017; Morgan *et al*., 2021). Parent-of-origin effects on endosperm development have also been seen in crosses between species with cellular-type endosperms. In *Mimulus* and *Solanum*, maternal-excess crosses seem to develop smaller endosperm cells that are rapidly degraded by the growing embryo, whereas paternal-excess crosses develop fewer, larger endosperm cells that produce bigger seeds (Roth *et al*., 2018, Coughlan *et al*., 2020). Together, these studies of endosperm development have begun to build a case for the importance of parental conflict in shaping effective ploidy. Our study builds on these earlier studies by finding a “smoking gun” – that is, a distinct region (i.e., the chalazal haustorium) that seems to be specifically targeted by parental conflict.

If there are different potential targets for parental conflict within a seed, why do we argue for the primacy of the chalazal haustorium? In species across the angiosperm phylogeny, this specialized region of the endosperm takes on diverse forms but invariably occurs at the maternal-filial boundary, where it often projects directly into maternal tissues (Povilus & Gehring, 2022). In *A. thaliana* and cereal crops (both with nuclear-type endosperm development), patterns of gene expression in chalazal tissues – or in analogous endosperm transfer cells – also point to their role in nutrient transfer, with upregulation of genes involved in sugar transport and metabolism (Thiel, 2014, Zhan *et al*., 2015, Picard *et al*., 2021). In addition to this direct role in nutrient acquisition, the *Arabidopsis* chalazal endosperm appears to exert indirect effects on the process by producing the signaling protein TERMINAL FLOWER1 (TFL1), which moves to the peripheral endosperm and initiates cellularization (Zhang *et al*., 2020). Thus, mounting evidence suggests genes expressed in the chalazal region are critical in determining the amount and timing of nutrient flow into the developing embryo.

Our finding that the chalazal endosperm develops abnormally in inviable, paternal-excess F1 hybrid *Mimulus* seeds also adds to a growing body of evidence suggesting this tissue is particularly sensitive to parental dosage and gene imprinting. Under a scenario of parental conflict in which maternally expressed genes (MEGs) and paternally expressed genes (PEGs) spar over the distribution of maternally-supplied resources to the developing seeds, the chalazal endosperm should play a key role (Povilus & Gehring, 2022). In line with this prediction, gene expression of two major regulators of PEGs in *A. thaliana* – *FIS2* and *MEA* – becomes localized in the chalazal cyst right at the point of cellularization (Luo *et al*., 2000). *FIS2* and *MEA* are themselves MEGs and members of the Polycomb Repressive Complex 2 (PRC2) complex, which act to epigenetically silence the maternal alleles of PEGs (Kinoshita *et al*., 1999; Luo *et al*., 2000; Köhler *et al*., 2005). In *fis2* mutants, endosperm cellularization fails, hexose accumulation in the central vacuole is prolonged (Hehenberger *et al*., 2012), and the chalazal endosperm is enlarged (sometimes filling ∼50% of the endosperm; Sørenson *et al*., 2001). This scenario of an evolutionary arms race between imprinted genes might explain why effective ploidy is positively correlated with the number and expression of PEGs in the endosperm of *Capsella* species (Lafon-Placette *et al*., 2018). Additionally, single nucleus RNA-sequencing in *Arabidopsis* shows that PEG expression is specifically enriched in the chalazal endosperm (Picard *et al*., 2021). Together with our study, this evidence points toward parental conflict driving rapid changes in gene expression within the chalazal endosperm because it is a particularly effective venue for manipulating the transfer of maternal resources. In further support of this idea, chalazal-specific genes in two species of *Arabidopsis* show elevated rates of adaptive evolution compared to genes expressed in other regions of the seed (Geist *et al*., 2019).

In addition to the chalazal haustorium, parental conflict might target other tissues in the developing seed that regulate nutrient transfer to the embryo, including the micropylar region, which transfers sucrose from the integuments to the embryo (Morley-Smith *et al*., 2008). We found that the micropylar haustorium typically degenerates before 10 DAP in intraspecific *Mimulus* crosses but persists in some paternal-excess crosses. For example, when *M. tilingii* acts as the seed parent and *M. guttatus* as the pollen parent, the micropylar region appears enlarged in developing hybrid seeds and is still present at 10 DAP (Fig S4). Similar, though less severe abnormalities also appear in CxT hybrid seeds, but a more detailed investigation of seed development in the micropylar region is needed. Intriguingly, disruptions to the micropylar region have also been reported in paternal-excess, interploidy crosses in *Galeopsis* and *Arabidopsis* (Håkansson, 1952; Scott *et al*., 1998), with micropylar haustoria vigorously invading seed integuments.

In addition to identifying the chalazal haustorium as a potential target of parental conflict, our study is one of only a handful to investigate divergence in effective ploidy among multiple, closely related species pairs. In this trio of *Mimulus* species, we find that effective ploidy is somewhat related to genetic distance – that is, the most closely related species pair, *M. caespitosa* and *M. tilingii*, has diverged the least in effective ploidy. However, the fact that each species has evolved to a different level of effective ploidy implies there have been lineage-specific changes, potentially driven by differences in the strength of parental conflict. The evolution of a relatively high effective ploidy in *M. guttatus* suggests that parental conflict has either increased in this species or decreased in the lineage leading to *M. caespitosa* and *M. tilingii*. Additionally, a lower effective ploidy in *M. caespitosa* might suggest this species has experienced a relaxation in conflict compared to *M. tilingii*. Consistent with this idea, *M. caespitosa* seems to have shifted toward self-fertilization, which theory predicts should decrease the opportunity for parental conflict (Brandvain & Haig, 2005). Although all three *Mimulus* species are hermaphroditic and self-compatible, *M. caespitosa* has a reduced anther-stigma distance and often self-fertilizes in the greenhouse (Sandstedt *et al*., 2021). The strength of parental conflict within species may also depend on other factors that influence effective population size (Coughlan *et al*., 2020, reviewed in Städler *et al*., 2021). In line with this expectation, nucleotide diversity in these three *Mimulus* species follows the same rank order as effective ploidy (Sandstedt *et al*., 2021). Even with these potentially divergent histories of conflict, disruption of the chalazal haustorium was observed in the F1 hybrid seeds of all *Mimulus* species pairs, which might suggest there have been parallel developmental changes across lineages. Going forward, identifying the genetic basis of these developmental phenotypes will be an important step toward understanding how and when parental conflict drives speciation.

## Acknowledgments

We thank Tylanna Baker and Jaylin Knight for help with data collection. We are grateful to Jill Anderson, Wolfgang Lukowitz, David Hall, Robert Schmitz, Robert Franks, Alex Sotola, Samuel Mantel, Matthew Farnitano, Makenzie Whitener, Jenn Coughlan, Elen Oneal, Miguel Flores-Vergara, Jay Sobel, and John Willis for helpful discussions. Robert Franks, Jenn Coughlan, Alex Sotola, Samuel Mantel, Elen Oneal, and John Willis provided valuable comments and improved the quality of the manuscript. This work was supported by the National Institutes of Health T32 Fellowship [GM007103] to G.D.S., the Jan and Kirby Alton Fellowship [Department of Genetics, UGA] to G.D.S., and National Science Foundation grants [DEB-1350935 and DEB-1856180] to A.L.S.

The authors declare no competing interests.

## Author Contributions

Research conceived and designed by G.D.S. and A.L.S., data collected and analyzed by G.D.S., and manuscript written by G.D.S. and A.L.S. G.D.S. and A.L.S. contributed equally.

## Data Accessibility

Data will be made available on Dryad Digital Repository.

## SUPPORTING INFORMATION

### Methods S1 Data analysis

**Table S1** The effect of intra- and interspecific crosses among *M. caespitosa* (C), *M. minor* (M), and *M. tilingii* (T) on the number of fully-developed seeds per fruit (scored by eye) as determined by generalized linear mixed models.

**Table S2** The effect of intra- and interspecific crosses among *M. caespitosa* (C), *M. minor* (M), and *M. tilingii* (T) on the number of seeds stained dark red by tetrazolium (*i*.*e*., viable seeds) as determined by generalized linear mixed models.

**Table S3** The effect of intra- and interspecific crosses among *M. caespitosa* (C), *M. minor* (M), and *M. tilingii* (T) on germination success using sucrose rich media as determined by generalized linear mixed models.

**Table S4** The effect of intra- and interspecific crosses among *M. caespitosa* (C), *M. minor* (M), and *M. tilingii* (T) on seed area as determined by linear mixed models.

**Table S5:** The effect of intra- and interspecific crosses among *M. caespitosa* (C), *M. minor* (M), and *M. tilingii* (T), days after pollination (DAP), and their interaction on the area of the endosperm filled by a chalazal haustorium as determined by linear models.

**Fig. S1** Tetrazolium assay for seed viability of intra-, interspecific, and interploidy crosses among *M. caespitosa* (C), *M. tilingii* (T), and *M. guttatus* (G).

**Fig. S2** Total seed area from crosses within and between *M. caespitosa* (C), *M. tilingii* (T), and *M. guttatus* (G).

**Fig. S3** Developing seeds 3 and 4 days after pollination (DAP) in crosses among *M. caespitose* (C), *M. tilingii* (T), and *M. guttatus* (G).

**Fig. S4** Replicate of Figure S3.3 without outlined structures.

**Fig. S5** Histological sections of whole fruits from intra- and interspecific crosses among *M. caespitosa* (C), *M. tilingii* (T), and *M. guttatus* (G) at 5, 6, 8, and 10 days after pollination (DAP).

## REFERENCES

Arekal GD. 1965. Embryology of Mimulus ringens. Botanical Gazette 126: 58–66.

Bates D, Sarkar D, Bates MD, Matrix L. 2007. The lme4 package. R package version 2: 74.

Batista RA, Köhler C. 2020. Genomic imprinting in plants-revisiting existing models. Genes & development 34: 24–36.

Baud S, Wuillème S, Lemoine R, Kronenberger J, Caboche M, Lepiniec L, Rochat C. 2005. The AtSUC5 sucrose transporter specifically expressed in the endosperm is involved in early seed development in Arabidopsis. Plant Journal 43: 824–836.

Berger F. 2003. Endosperm: the crossroad of seed development. Current opinion in plant biology 6: 42–50

Berger F, Hamamura Y, Ingouff M, Higashiyama T. 2008. Double fertilization – caught in the act. Trends in Plant Science 13: 437–443.

Brandvain Y, Haig D. 2005. Divergent Mating Systems and Parental Conflict as a Barrier to Hybridization in Flowering Plants. The American Naturalist 166: 30–338.

Brink RA, Cooper DC. 1947. The Endosperm in Seed Development. The Botanical Review 13: 479–541.

Brown RC, Lemmon BE, Nguyen H. 2003. Events during the first four rounds of mitosis establish three developmental domains in the syncytial endosperm of Arabidopsis thaliana. Protoplasma 222: 167–174.

Bushell C, Spielman M, Scott RJ. 2003. The basis of natural and artificial postzygotic hybridization barriers in Arabidopsis species. Plant Cell 15: 1430–1442.

Cooper DC, and Brink RA. 1942. The endosperm as a barrier to interspecific hybridization in flowering plants. Science 95: 75–76.

Coughlan JM, Brown MW, Willis JH. 2020. Patterns of Hybrid Seed Inviability in the Mimulus guttatus sp. Complex Reveal a Potential Role of Parental Conflict in Reproductive Isolation. Current Biology 30: 83–93.

Coughlan, JM, Brown MW, and Willis JH. 2021. The genetic architecture and evolution of life-history divergence among perennials in the Mimulus guttatus species complex. Proceedings of the Royal Society B. 288: p. 20210077.

Dobzhansky T. 1937. Genetic nature of species differences. The American Naturalist. 71: 404–420.

Floyd SK, Friedman WE. 2000. Evolution of endosperm developmental patterns among basal flowering plants. International Journal of Plant Sciences 161: S57–S81.

Garcia D, Saingery V, Chambrier P, Mayer U, Jürgens G, Berger F. 2003. Arabidopsis haiku mutants reveal new controls of seed size by endosperm. Plant Physiology 131: 1661–1670.

Garner AG, Kenney AM, Fishman L, Sweigart, AL. 2016. Genetic loci with parent‐of‐origin effects cause hybrid seed lethality in crosses between Mimulus species. New Phytologist 211: 319–331.

Geist KS, Strassmann JE, Queller DC. 2019. Family quarrels in seeds and rapid adaptive evolution in Arabidopsis. Proceedings of the National Academy of Sciences 116: 9463–9468

Guilford VB, Fisk EL. 2016. Torrey Botanical Society Megasporogenesis and Seed Development in Mimulus tigrinus and Torenia fournieri. Bulletin of the Torrey Botanical Club. 79: 6–24.

Haig D, Westoby M. 1989. Parent-Specific Gene Expression and the Triploid Endosperm. The American Naturalist 134: 147–155.

Haig D, Westoby M. 1991. Genomic Imprinting in Endosperm: Its Effect on Seed Development in Crosses between Species, and between Different Ploidies of the Same Species, and Its Implications for the Evolution of Apomixis. Philosophical Transactions: Biological Sciences 1–13.

Håkansson A. 1952. Seed development after 2x, 4x crosses in Galeopsis pubescens. Hereditas 38:425–448.

Hamilton WD. 1964. The genetical theory of kin selection. J. Theor. Biol, 7: 1–52.

Hehenberger E, Kradolfer D, Köhler C. 2012. Endosperm cellularization defines an important developmental transition for embryo development. Development 139: 2031–2039.

İltaş Ö, Svitok M, Cornille A, Schmickl R, Lafon Placette C. 2021. Early evolution of reproductive isolation: A case of weak inbreeder/strong outbreeder leads to an intraspecific hybridization barrier in Arabidopsis lyrata. Evolution 75: 1466–1476.

Johnston SA, den Nijs TPM, Peloquin SJ, Hanneman RE. 1980. The Significance of Genic Balance to Endosperm Development in Interspecific Crosses. Theoretical and applied genetics 57: 5–9.

Kang IH, Steffen JG, Portereiko MF, Lloyd A, Drews GN. 2008. The AGL62 MADS domain protein regulates cellularization during endosperm development in Arabidopsis. Plant Cell 20: 635–647.

Kinoshita T, Yadegari R, Harada JJ, Goldberg RB, and Fischer RL. 1999. Imprinting of the MEDEA polycomb gene in the Arabidopsis endosperm. The Plant Cell 11: 1945–1952.

Kinoshita T. 2007. Reproductive barrier and genomic imprinting in the endosperm of flowering plants. Genes & genetic systems 82: 177–186.

Kinser TJ, Smith RD, Lawrence AH, Cooley AM, Vallejo-Marín M, Conradi Smith GD, Puzey JR. 2021. Endosperm-based incompatibilities in hybrid monkeyflowers. The Plant Cell 33: 2235–2257.

Köhler C., Page DR, Gagliardini V, and Grossniklaus U. 2005. The Arabidopsis thaliana MEDEA Polycomb group protein controls expression of PHERES1 by parental imprinting. Nature genetics 37: 28–30.

Lafon-Placette C, Hatorangan MR, Steige KA, Cornille A, Lascoux M, Slotte T, Köhler C. 2018. Paternally expressed imprinted genes associate with hybridization barriers in Capsella. Nature Plants 4: 352–357.

Lafon-Placette C, Johannessen IM, Hornslien KS, Ali MF, Bjerkan KN, Bramsiepe J, Glöckle BM, Rebernig CA, Brysting AK, Grini PE, Köhler C. 2017. Endosperm-based hybridization barriers explain the pattern of gene flow between Arabidopsis lyrata and Arabidopsis arenosa in Central Europe. Proceedings of the National Academy of Sciences of the United States of America 114: E1027–E1035.

Lafon-Placette C, Köhler C. 2016. Endosperm-based postzygotic hybridization barriers: developmental mechanisms and evolutionary drivers. Molecular ecology 25: 2620–2629.

Lenth R, Lenth MR. 2018. Package ‘lsmeans’. The American Statistician, 34: 216–221.

Lin B-Y. 1984. Ploidy barrier to endosperm development in maize. Genetics 107: 103–115.

Lu J, Zhang C, Baulcombe DC, Chen ZJ. 2012. Maternal siRNAs as regulators of parental genome imbalance and gene expression in endosperm of Arabidopsis seeds. Proceedings of the National Academy of Sciences of the United States of America 109: 5529–5534.

Lu Z, Hofmeister BT, Vollmers C, DuBois RM, and Schmitz RJ. 2017. Combining ATAC- seq with nuclei sorting for discovery of cis-regulatory regions in plant genomes. Nucleic acids research, 45: 41–41.

Luo M, Bilodeau P, Dennis ES, Peacock WJ, Chaudhury A. 2000. Expression and parent-of- origin effects for FIS2, MEA, and FIE in the endosperm and embryo of developing Arabidopsis seeds. Proceedings of the National Academy of Sciences 97: 10637–10642.

Luo M, Dennis ES, Berger F, Peacock WJ, Chaudhury A. 2005. MINISEED3 (MINI3), a WRKY family gene, and HAIKU2 (IKU2), a leucine-rich repeat (LRR) KINASE gene, are regulators of seed size in Arabidopsis. Proceedings of the National Academy of Sciences 102: 17531–17536.

Mikesell J. 1990. Anatomy of terminal haustoria in the ovule of plantain (Plantago major L.) with taxonomic comparison to other angiosperm taxa. Botanical Gazette, 151: 452–464.

Morgan EJ, Čertner M, Lučanová M, Deniz U, Kubíková K, Venon A, Kovářík O, Lafon Placette C, Kolář, F. 2021. Disentangling the components of triploid block and its fitness consequences in natural diploid–tetraploid contact zones of Arabidopsis arenosa. New Phytologist 232: 449–1462.

Morley-Smith ER, Pike MJ, Findlay K, Köckenberger W, Hill LM, Smith AM, Rawsthorne S. 2008. The transport of sugars to developing embryos is not via the bulk endosperm in oilseed rape seeds. Plant Physiology 147: 2121–2130.

Muller H. J. 1942. Isolating mechanisms, evolution, and temperature. Biological Symposium 6: 71–125.

Nesom GL. 2012. Taxonomy of Erythranthe sect. Simiola (Phrymaceae) in the USA and Mexico. Phytoneuron 40: 1–123.

Nguyen H, Brown RC, Lemmon, BE. 2000. The specialized chalazal endosperm in Arabidopsis thaliana and Lepidium virginicum (Brassicaceae). Protoplasma, 212: 99–110.

Nishiyama I, Inomata N. 1966. Embryological studies on cross-incompatibility between 2x and 4x in Brassica. The Japanese journal of genetics 41: 27–42.

Nishiyama I, Yabuno T. 1978. Causal Relationships between the Polar Nuclei in Double Fertilization and Interspecific Cross-incompatibility in Avena. Cytologia, 43: 453–466.

Oneal E, Willis JH, Franks RG. 2016. Disruption of endosperm development is a major cause of hybrid seed inviability between Mimulus guttatus and Mimulus nudatus. New Phytologist 210: 1107–1120.

Pennington PD, Costa LM, Gutierrez-Marcos JF, Greenland AJ, Dickinson HG. 2008. When genomes collide: Aberrant seed development following maize interploidy crosses. Annals of botany 101: 833–843.

Picard CL, Povilus RA, Williams BP, Gehring M. 2021. Transcriptional and imprinting complexity in Arabidopsis seeds at single-nucleus resolution. Nature plants 7: 730–738.

Povilus RA, Gehring M. 2022. Maternal-filial transfer structures in endosperm: A nexus of nutritional dynamics and seed development. Current opinion in plant biology 65: 102121.

Queller DC. 1983. Kin Selection and Conflict in Seed Maturation. Journal of Theoretical Biology 100: 153–172.

Rasband WS. 1997. ImageJ, U. S. National Institutes of Health, Bethesda, Maryland, USA, https://imagej.nih.gov/ij/.

Rebernig CA, Lafon-Placette C, Hatorangan MR, Slotte T, Köhler C. 2015. Non-reciprocal interspecies hybridization barriers in the Capsella genus are established in the endosperm. PLoS genetics 11:1005295.

Reik W, Walter J. 2001. Genomic imprinting: parental influence on the genome. Nature Reviews Genetics 2: 21–32.

Roth M, Florez-Rueda AM, Griesser S, Paris M, Städler T. 2018. Incidence and developmental timing of endosperm failure in post-zygotic isolation between wild tomato lineages. Annals of Botany 121: 107–118.

Sandstedt GD, Wu CA, Sweigart AL. 2021. Evolution of multiple postzygotic barriers between species of the Mimulus tilingii complex*. Evolution 75: 600–613.

Scott RJ, Spielman M, Bailey J, Dickinson HG. 1998. Parent-of-origin effects on seed development in Arabidopsis thaliana. Development 125: 3329–3341.

Sørensen MB, Chaudhury AM, Robert H, Banchare E, Berger F. 2001. Polycomb group genes control pattern formation in plant seed. Current Biology 11: 277–281.

Städler T, Florez-Rueda AM, Roth M. 2021. A revival of effective ploidy: the asymmetry of parental roles in endosperm-based hybridization barriers. Current Opinion in Plant Biology 61: 102015

Stephens SG. 1949. The cytogenetics of speciation in Gossypium. I. Selective elimination of the donor parent genotype in interspecific backcrosses. Genetics 34: 627.

Thiel J. 2014. Development of endosperm transfer cells in barley. Frontiers in Plant Science 5: 108.

Vickery RK. 1978. Case studies in the evolution of species complexes in Mimulus. Evolutionary biology 405–507. Springer, Boston, MA.

Woodell SRJ, Valentine DH. 1961. Studies in British Primulas IX. Seed Incompatibility in Diploid-Autotetraploid Crosses. New Phytologist 282–294.

Zhan J, Thakare D, Ma C, Lloyd A, Nixon NM, Arakaki AM, Burnett WJ, Logan KO, Wang D, Wang X, Drews GN. 2015. RNA sequencing of laser-capture microdissected compartments of the maize kernel identifies regulatory modules associated with endosperm cell differentiation. The Plant Cell, 27: 513–531.

Zhang B, Li C, Li Y, and Yu, H. 2020. Mobile TERMINAL FLOWER1 determines seed size in Arabidopsis. Nature Plants 6: 1146–1157.

