## Supplementary material for "Developmental evidence for parental conflict in driving *Mimulus* species barriers": Fig. S1

**Fig. S1** Tetrazolium assay for seed viability of intra-, interspecific, and interploidy crosses among *M. caespitosa* (C), *M. tilingii* (T)*,* and *M. guttatus* (G). **(a)** Example of tetrazolium test on seeds from intra and interspecific crosses of *M. tilingii* and *M. caespitosa.* Intraspecific crosses: TxT (top left) and CxC (bottom right). Interspecific crosses, maternal parent is always listed first: CxT (bottom left), TxC (top right). Dark red seeds are scored as viable, and pink or white seeds are scored as inviable. Scale bar is 1 mm. **(b)** Percentage of seeds stained red from intra- and interspecific crosses. Least squares means (lsmeans) given with +/- SE. Light gray bars represent crosses between diploid parents, and dark gray bars represent crosses where one parent is a synthetic tetraploid, as denoted by the “4n” subscript in the cross. Different letters indicate significant differences in lsmeans among crosses (P<0.05) determined by a post hoc Tukey method. Analyses were performed separately, only comparing reciprocal interspecific and interploidy crosses and their corresponding intraspecific crosses. Asterisk denotes insufficient variation in response variable to determine statistical differences. Note that for some interploidy crosses, 5-10 fully-developed seeds were planted to test for ploidy prior to tetrazolium assay.

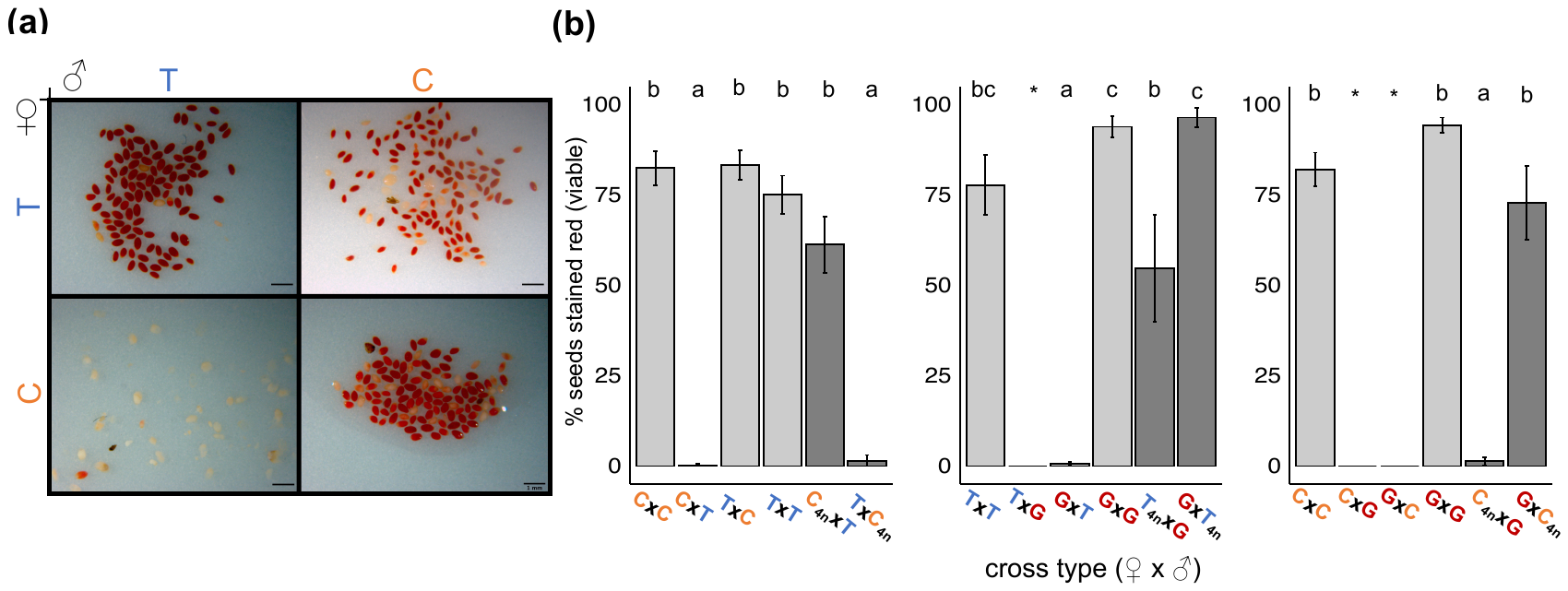
