## Supplementary material for "Developmental evidence for parental conflict in driving *Mimulus* species barriers": Fig. S2

**Fig. S2** Total seed area from crosses within and between *M. caespitosa* (C), *M. tilingii* (T)*,* and *M. guttatus* (G). The first letter of each cross indicates the maternal species. Different letters indicate significant differences in least-squares means among crosses (P<0.05) determined by a post hoc Tukey method. Analyses were performed separately, only comparing reciprocal interspecific crosses and their corresponding intraspecific crosses

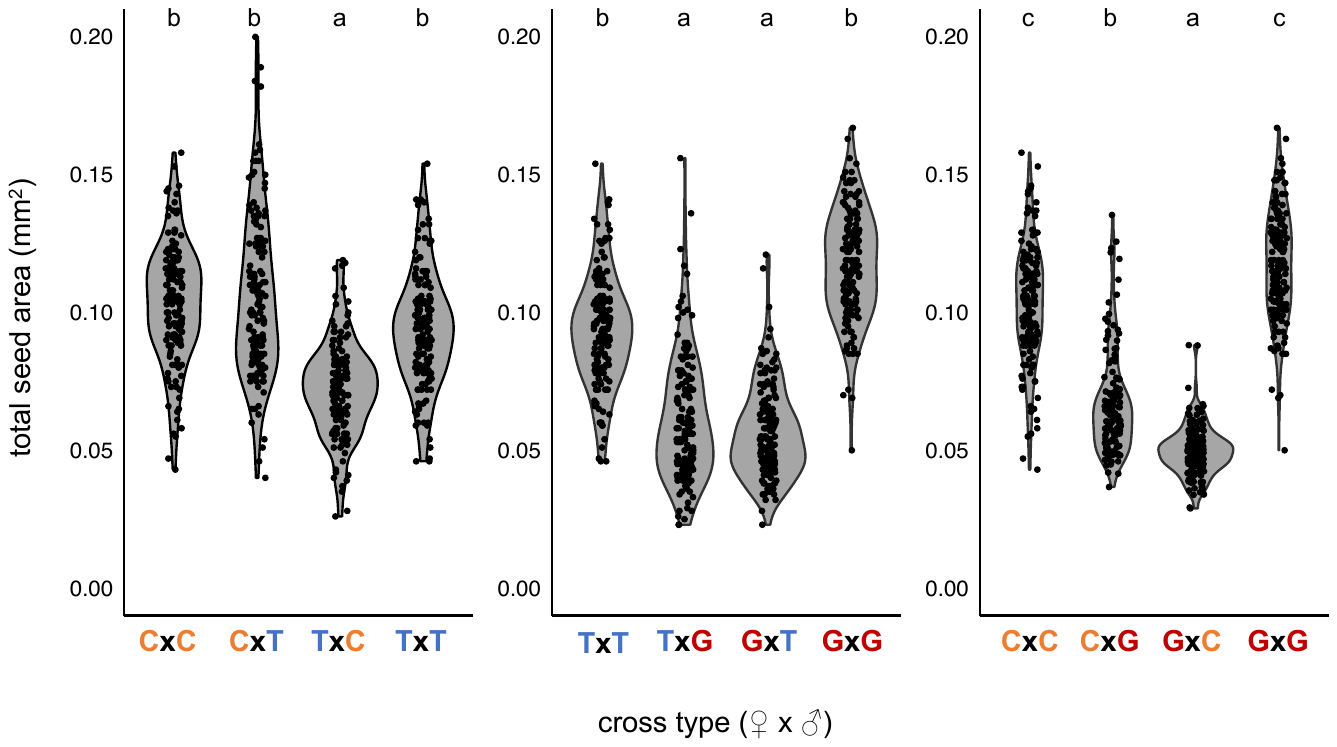
