## Supplementary material for "Developmental evidence for parental conflict in driving *Mimulus* species barriers": Fig. S4

**Fig. S4** Replicate of Figure S3.3 without outlined structures. Developing seeds 3 and 4 days after pollination (DAP) in crosses among *M. caespitosa* (C), *M. tilingii* (T)*,* and *M. guttatus* (G). We also included one interploidy cross (T_4N_xG). Maternal parent is listed first in interspecific crosses. Seeds were cleared with Hoyer’s solution. Scale bar is 0.1mm. At 3 DAP, chalazal and micropylar haustoria domains are fully established. The micropylar domain is composed of two cells (arrow points to two nuclei in micropylar region of the TxT seed) at the anterior end of the seed, and this region invades nearby seed integuments. We also sometimes observe micropylar haustorium extending towards the chalazal domain (see GxG, GxT, and GxC). The chalazal haustorium is composed of two cells that occupy the posterior end of the seed. The chalazal haustorium extends from the maternal-filial boundary towards the anterior end of the seed. At 4 DAP, the chalazal haustorium has largely degenerated in TxT, TxC, GxT, GxC, and T_4N_xG crosses, as the central endosperm proliferates. The area of the endosperm that is filled by the chalazal haustorium decreases from 3 to 4 DAP in almost all crosses, except for CxG

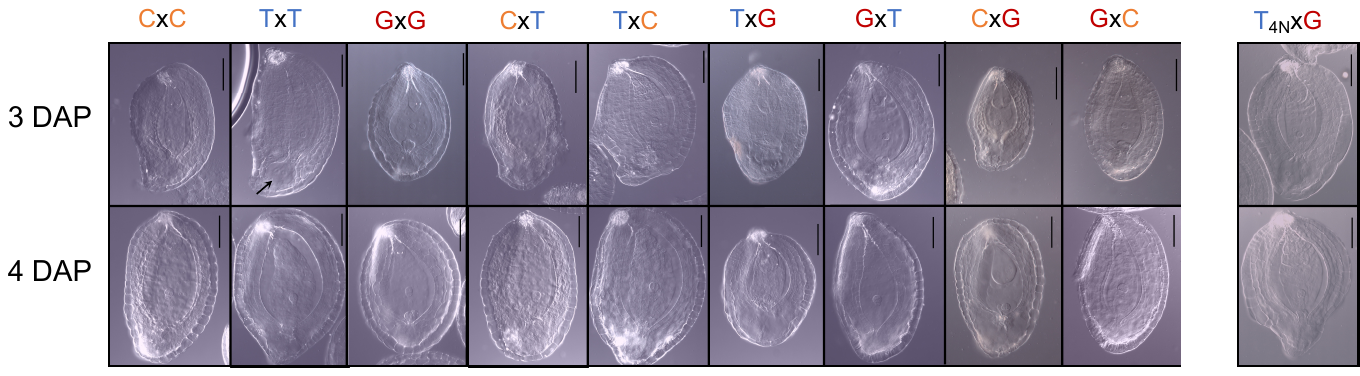
