## Supplementary material for "Developmental evidence for parental conflict in driving *Mimulus* species barriers": Fig. S5

**Fig. S5** Histological sections of whole fruits from intra- and interspecific crosses among *M. caespitosa* (C), *M. tilingii* (T)*,* and *M. guttatus* (G) at 5, 6, 8, and 10 days after pollination (DAP). Maternal parent is listed first in all interspecific crosses. Arrowhead = embryo, en = endosperm, sc = seed coat, ch = chalazal haustorium, mh = micropylar haustorium. Note that ch and mh are only labeled when haustoria are visible in the image. Scale bar is 0.1mm. At **5 and 6 DAP**, intraspecific crosses have reached a globular embryo stage, where the embryo is surrounded by ‘empty’ cells, and the chalazal haustorium has degenerated. Embryos of GxT and GxC maternal excess crosses are surrounded by dense, starch-filled cells, again with no chalazal haustorium present. In paternal-excess crosses (CxT, TxG, and CxG), embryos have not yet reached a full globular stage, the chalazal haustorium is still intact in some seeds of paternal-excess crosses at 5 DAP and in TxG and CxG at 6 DAP. We also note here that chalazal haustorium of CxT and TxG are deeply stained, likely with sugars, while the CxG haustorium is large and unstained. At **8DAP**, intraspecific and maternal-excess crosses have reached the heart shaped embryo stage, though heart embryos of maternal-excess crosses GxT and GxC appear abnormal. While in the intraspecific crosses, the central endosperm cells begin to break down and the peripheral endosperm near the seed coat starts to differentiate into starch-filled cells, the maternal-excess crosses appear fully differentiated and the endosperm area appears reduced. In contrast, endosperm cells in paternal-excess crosses remain empty and enlarged, and embryos are underdeveloped. By **10 DAP**, all intraspecific crosses and TxC have developed torpedo shaped embryos surrounded by a few layers of dense, starch-filled cells, and micropylar haustoria are completely degenerated. The maternal-excess crosses, GxT and GxC, fail to develop torpedo shaped embryos and remain as abnormal heart shaped embryos, with little to no endosperm and no apparent micropylar haustorium. In CxT, the embryo has finally reached a heart shape, but the endosperm cells remain undifferentiated and micropylar cells are evident in some seeds. While TxG seeds are severely underdeveloped, with prominent micropylar haustorium in some seeds, the seeds of CxG crosses have already collapsed around the underdeveloped embryo.
