## Supplementary material for "Developmental evidence for parental conflict in driving *Mimulus* species barriers": Supp. Methods 1

**SUPPORTING METHODS**

**Methods 1** ***Data Analysis***

We modeled the effect of cross on seed area using three separate linear mixed models, each with four comparisons including reciprocal interspecific crosses and the corresponding intraspecific crosses (CxC, CxT, TxC, TxT; TxT, TxG, GxT, GxG; and CxC, CxG, GxC, GxG). For each model, we fit a Gaussian distribution using the lmer command in the “lme4” package implemented in R (Bates *et al.*, 2007). We assigned our fixed factor as cross, random factor as individual plant, and our response variable as seed area (mm^2^). To determine whether there was an effect of cross on the variance of seed area, we computed an ANOVA test using the anova function in the R package “car” with type III sums of squares, which applies Wald chi-square tests for mixed models. We calculated least-squares means (lsmeans) using the emmeans function in the R package “emmeans”, performed pairwise comparisons between all crosses, and we used a post hoc Tukey method adjustment to determine which crosses differed significantly in seed area (Lenth & Lenth, 2018).

We determined the effect of cross on seed viability using three generalized linear mixed models (GLMMs), for both measures of seed viability (visual and tetrazolium assessment). Each GLMM compared reciprocal interspecific crosses, and their corresponding interploidy and intraspecific crosses (CxC, CxT, TxC, TxT, C_4n_xT, TxC_4n_; TxT, TxG, GxT, GxG, T_4n_xG, GxT_4n_; and CxC, CxG, GxC, GxG, C_4n_xG, GxC_4n_). In these models, we fit GLMMs with a binomial distribution using the glmer command in the “lme4” package implemented in R (Bates *et al.*, 2007). For our response variable, we combined the number of viable seeds (fully-developed or stained dark red) and the number of inviable seeds (under-developed or unstained) into a single variable using the R function cbind. We assigned our fixed factor as cross, and the individual plant was set as a random factor. We computed ANOVAs using the anova function to determine whether cross significantly affected the variance of seed viability. Then, we calculated lsmeans and performed pairwise comparisons between all crosses. We determined which crosses differed significantly in the number of viable seeds using a post hoc Tukey method adjustment.

To model the effect of cross on germination success of seed viability rescues with sucrose media, we performed three separate GLMMs, comparing only reciprocal interspecific crosses and their corresponding intraspecific crosses (CxC, CxT, TxC, TxT; TxT, TxG, GxT, GxG; and CxC, CxG, GxC, GxG). In these models, we fit GLMMs with a binomial distribution using the glmer command. For our response variable, we combined the number of seeds that germinated and the number of seeds that failed to germinate on a sucrose-rich medium into a single variable using the R function cbind. We assigned our fixed factor as cross, and the individual plant was set as a random factor. We computed an ANOVA to determine which crosses significantly affected variance of germination success on a sucrose-rich medium using the anova function in R. Similar to prior analyses, we estimated lsmeans, performed pairwise comparisons of lsmeans between all crosses, and determined which crosses significantly differed in the number of seeds that germinated on a sucrose-rich medium using a post hoc Tukey method.

To determine whether cross had a significant effect on area of endosperm filled by a chalazal haustorium, we performed three separate linear models for both measurements, comparing only reciprocal interspecific crosses and their corresponding intraspecific crosses—except for T-G comparisons, in which case we also included measurements of the interploidy cross (CxC, CxT, TxC, TxT; TxT, TxG, GxT, GxG, T_4n_xG; and CxC, CxG, GxC, GxG). We fit linear models using the lm function in R, assigning the response variable as either chalazal haustorium/endosperm area and fixed factors as cross, DAP, and their interaction. To determine whether these fixed factors affected the variance of the response variables, we computed ANOVAs with type III sums of squares. Then, we estimated lsmeans, performed pairwise comparisons of lsmeans, and determined which crosses at 3 and 4 DAP differed in embryo area and area of the endosperm filled by the chalazal haustorium.
