## Supplementary material for "Developmental evidence for parental conflict in driving *Mimulus* species barriers": Table S1

**Table S1** The effect of intra- and interspecific crosses among *M. caespitosa* (C), *M. minor* (M), and *M. tilingii* (T) on the number of fully-developed seeds per fruit (scored by eye) as determined by generalized linear mixed models. In these models, we also included interspecific, interploidy crosses (as denoted by 4-2 and 2-4, with the first number indicating the maternal parent’s ploidy level). For all interspecific crosses, maternal parent is always listed first. Three separate models were performed, each with six comparisons including reciprocal interspecific crosses, interspecific interploidy crosses, and their corresponding intraspecific crosses (CxC, CxT, CxT 4-2, TxC, TxC 2-4, TxT; TxT, TxG, TxG 4-2, GxT, GxT 2-4, GxG; and CxC, CxG, CxG 4-2, GxC, GxC 2-4, GxG). On the left, output from an ANOVA, with type III sums of squares, chi-square (χ2), degrees of freedom (df), and *P*-values calculated using likelihood ratio tests. On the right, least-squares means (lsmeans) and standard error (SE) for each cross. Lsmeans denoted by a different letter (under “group”) indicates significant differences among crosses (*P <*0.05) determined by post-hoc Tukey method.

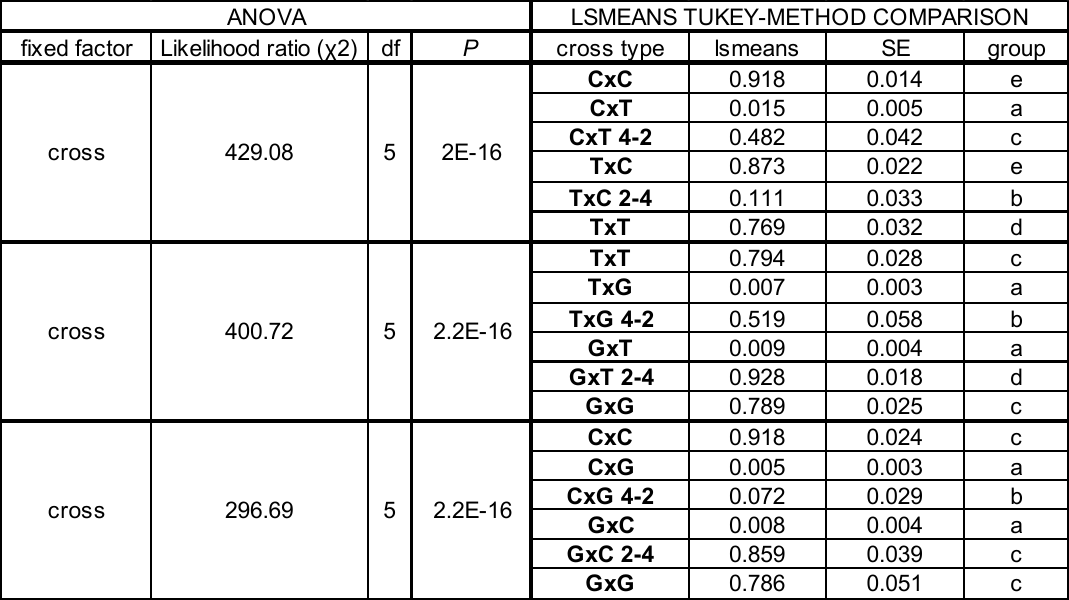
