## Supplementary material for "Developmental evidence for parental conflict in driving *Mimulus* species barriers": Table S3

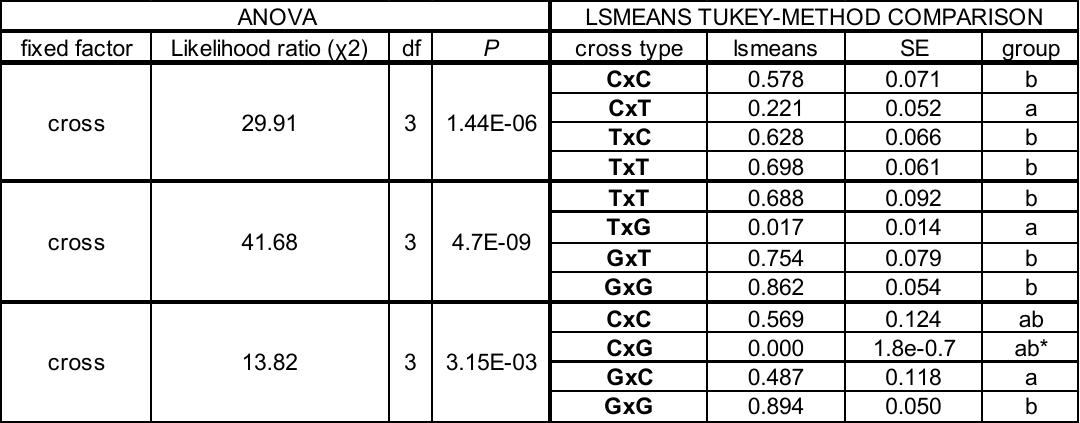
**Table S3** The effect of intra- and interspecific crosses among *M. caespitosa* (C), *M. minor* (M), and *M. tilingii* (T) on germination success using sucrose rich media as determined by generalized linear mixed models. For all interspecific crosses, maternal parent is always listed first. Three separate models were performed, each with four comparisons including reciprocal interspecific crosses and their corresponding intraspecific crosses (CxC, CxT, TxC, TxT; TxT, TxG, GxT, GxG; and CxC, CxG, GxC, GxG). On the left, output from an ANOVA, with type III sums of squares, chi-square (χ2), degrees of freedom (df), and *P*-values calculated using likelihood ratio tests. On the right, least-squares means (lsmeans) and standard error (SE) for each cross. Lsmeans denoted by a different letter (under “group”) indicates significant differences among crosses (*P <*0.05) determined by post-hoc Tukey method. Asterisk under “group” denotes insufficient variation in response variable to determine statistical differences.
