## Supplementary material for "Developmental evidence for parental conflict in driving *Mimulus* species barriers": Table S4

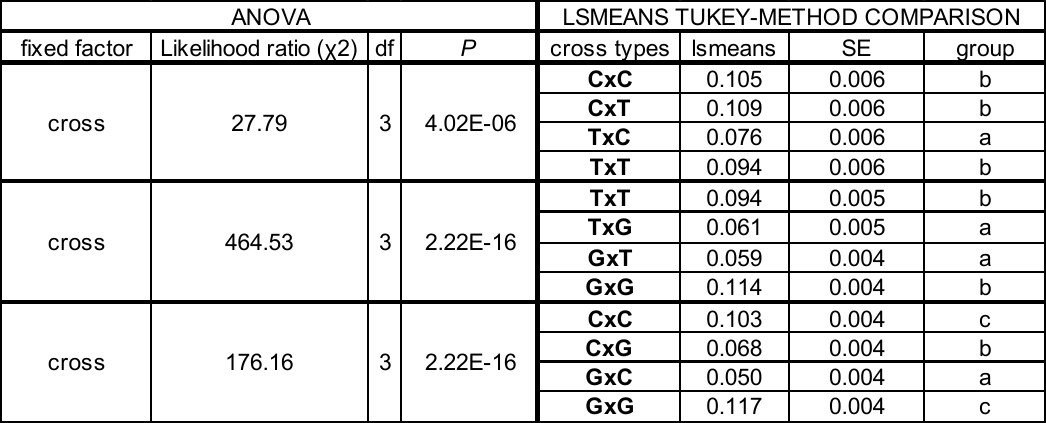
**Table S4** The effect of intra- and interspecific crosses among *M. caespitosa* (C), *M. minor* (M), and *M. tilingii* (T) on seed area as determined by linear mixed models. For all interspecific crosses, maternal parent is always listed first. Three separate models were performed, each with four comparisons including reciprocal interspecific and their corresponding intraspecific crosses (CxC, CxT, TxC, TxT; TxT, TxG, GxT, GxG; and CxC, CxG, GxC, GxG). On the left, output from an ANOVA, with type III sums of squares, chi-square (χ2), degrees of freedom (df), and *P*-values calculated using likelihood ratio tests. On the right, least-squares means (lsmeans) and standard error (SE) for each cross. Lsmeans denoted by a different letter (under “group”) indicates significant differences among crosses (*P <*0.05) determined by post-hoc Tukey method.
