## Supplementary material for "Developmental evidence for parental conflict in driving *Mimulus* species barriers": Table S5

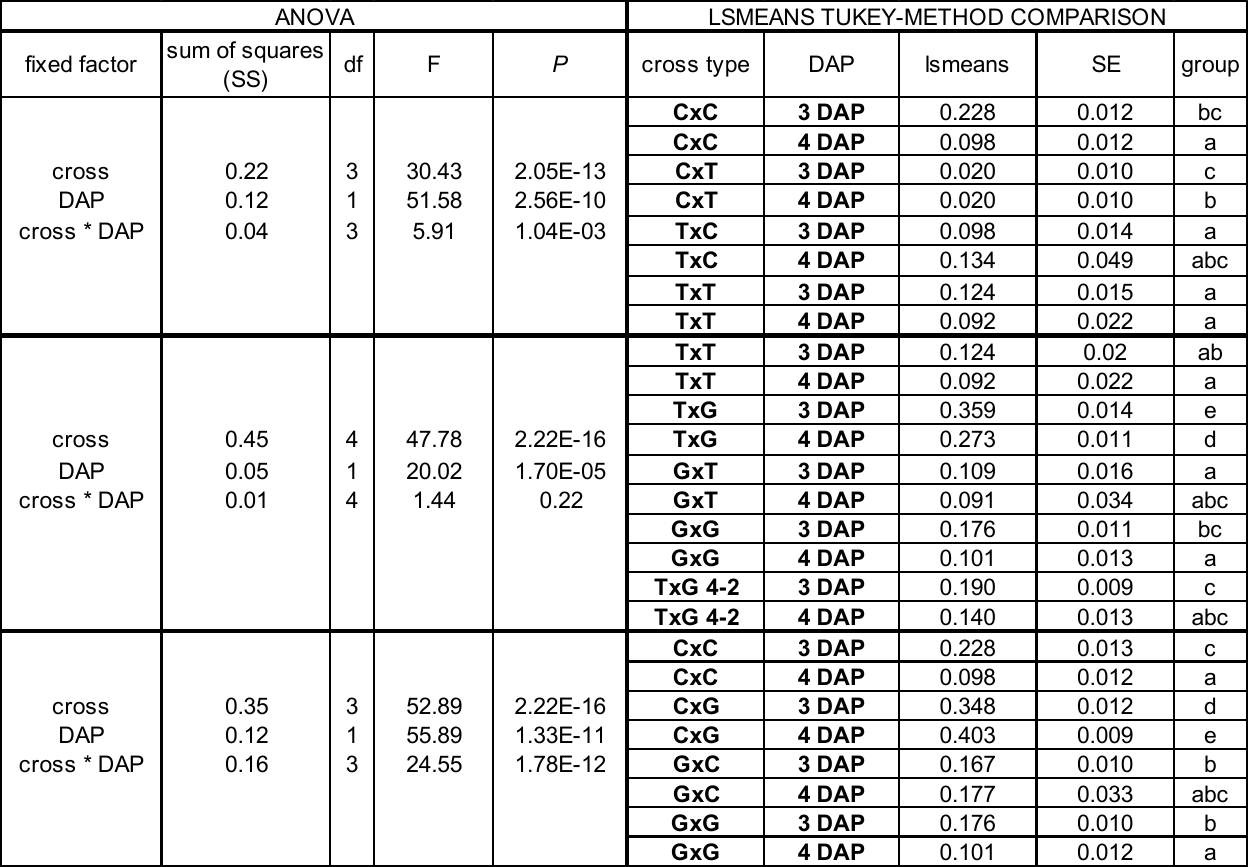
**Table S5** The effect of intra- and interspecific crosses among *M. caespitosa* (C), *M. minor* (M), and *M. tilingii* (T), days after pollination (DAP), and their interaction on the area of the endosperm filled by a chalazal haustorium as determined by linear models. For all interspecific crosses, maternal parent is always listed first. Three separate models were performed, two models had eight comparisons, including reciprocal interspecific crosses and their corresponding intraspecific crosses (3 and 4 DAP: CxC, CxT, TxC, TxT; CxC, CxG, GxC, GxG). One model had 10 comparisons, including reciprocal interspecific crosses, one interploidy cross with a tetraploid maternal and diploid paternal parent, and their corresponding intraspecific crosses (3 and 4 DAP: TxT; TxT, TxG, TxG 4-2, GxT, GxG). On the left, output from an ANOVA with type III sums of squares, sums of squares (SS), degrees of freedom (df), and *P*-values. On the right, least-squares means (lsmeans) and standard error (SE) for each cross. Lsmeans denoted by a different letter (under “group”) indicates significant differences among crosses (*P <*0.05) determined by post-hoc Tukey method.
